# Pancreatic Cancer Induces Population-Specific Switching of Myosin Isoforms and Discrete Activation of Cachexia Genes in Skeletal Muscle Myocytes

**DOI:** 10.1101/2025.09.01.673500

**Authors:** Brittany R. Counts, Sephora Jean, Omnia Gaafer, Sara K. Ota, Iishaan Inabathini, Tingbo Guo, Denis C. Guttridge, Michael C. Ostrowski, Leonidas G. Koniaris, Sha Cao, Hyun C. Roh, Teresa A. Zimmers

## Abstract

Skeletal muscle loss in pancreatic cancer is a significant cause of morbidity and mortality for patients. In order to understand myocytes changes we examined myonuclei- and myofiber-specific dynamics during pancreatic cancer cachexia progression. Single-nucleus RNA-seq was used to interrogate myonuclear gene expression, and RNAscope and immunofluorescence characterized myofiber-specific changes. Bulk RNA-seq of skeletal muscle provided a whole-muscle transcriptomic profile. Cachexia induces a progressive loss of muscle differentiation factor *Maf* and its target *Myh4*, accompanied by increased expression of *Myh1* and *Myh2*. This myofiber dedifferentiation occurs without evidence for fiber type shifting, regeneration, or proliferation. Single-nuclei analysis reveals global shifts in myofiber gene expression identity including the identification of a cachexia only myonuclear subpopulation. Cachexia gene expression was not restricted solely to this PDAC-specific myonuclear subpopulation and did not overlap with *Myh1* and *Myh2* expressing myonuclei early in cachexia. Altogether, PDAC cachexia elicits distinct transcriptional responses across different myonuclear populations. These results reveal population-specific heterogeneity in cachexia gene activation, rather than a uniform upregulation of cachexia mediators across muscle tissue. Our data suggest that myonuclei fate occurs prior to overt muscle wasting when cachexia gene expression only modestly overlaps with differentiation factors, with a strong association after irreversible muscle wasting. These findings explain the challenge of effectively targeting skeletal muscle wasting in cancer cachexia requires addressing the changing cell population induced through non overlapping mechanisms.

## Introduction

Cancer cachexia leads to systemic metabolic dysfunction with severe muscle loss and weakness leading to loss of mobility, increased frailty, and death. Pancreatic ductal adenocarcinoma (PDAC) accounts for most pancreatic cancer diagnoses and has a 5-year survival rate of 13%[1]. Cachexia in pancreatic cancer is severe with 60-70% of PDAC patients presenting with cachexia at diagnosis[2–4]. The hallmark, and most important functional consequence of cachexia, is the wasting of skeletal muscle and associated muscle weakness which can occur months prior to diagnosis in PDAC[5]. Identifying strategies to mitigate and prevent skeletal muscle wasting is imperative for treating cancer cachexia and improving patient survival.

Our understanding of the deleterious transcriptional and proteomic changes in skeletal muscle during cancer cachexia has traditionally been limited to analyses at the whole tissue level. Muscle fibers are comprised mostly of skeletal muscle tissue; it is often assumed that observed changes reflect alterations in muscle cell specificity. However, advances in single-cell technologies now allow us to distinguish which cell types are driving changes in response to muscle wasting. Our recent work has identified inflammatory cell activity promoting the wasting of muscle tissue, emphasizing the importance of cell type perturbations to wasting[6]. Despite the promise of single-cell RNA sequencing, its application in skeletal muscle is limited by the tissue’s unique characteristics; the large myofiber size, their multinucleated structure, and the fresh tissue requirement, which pose challenges for large-scale studies. These limitations underscore the need for single-nucleus RNA sequencing (snRNAseq), which overcomes many of these challenges and is increasingly applied to skeletal muscle research[7, 8].

Extensive research has demonstrated that cancer cachexia disrupts skeletal muscle protein homeostasis by suppressing protein synthesis and promoting a chronic catabolic state. This state is marked by increased inflammatory signaling, autophagy, and proteasome activity; collectively referred to as the molecular cachexia signature. Key pathways include: IL6/Gp130/Stat3[9, 10], NF-κB[11, 12], MAPK[13], and heat shock protein signaling[14], all of which are sufficient to induce muscle wasting. Although these catabolic conditions are known to be transcriptionally upregulated during cachexia progression[15, 16], it is unclear whether such transcriptional changes occur specifically within myonuclei rather than in other cell types within the microenvironment. This distinction is highly significant, as the activation of a given gene in myonuclei may have profoundly different implications compared to its activation in immune or endothelial cells. While muscle cross-section analysis illustrates fiber-specific regulation of cachexia-associated mediators[6, 17, 18], we often assume that myonuclei within a single fiber respond uniformly. However, the extent to which cachexia differentially regulates individual myonuclei within the same fiber remains unknown thus highlighting a critical gap in our understanding of intrafiber heterogeneity during muscle wasting.

Recent snRNAseq studies in KIC autochthonous pancreatic cancer and LLC syngeneic lung cancer cachexia mouse models have identified distinct catabolic myonuclei populations[19, 20]. These studies were primarily exploratory and conducted in an advanced stage of muscle wasting but lacked myofiber dynamics and validation in human tissue. To address this, we examined myonuclear alterations in a PDAC cachexia model before overt muscle loss and in patients with PDAC. Our analysis revealed early disruption of myofiber transcriptional identity including heterogeneous expression of myosin heavy chains and the progressive loss of the myocyte identity factor, Maf, and its downstream target Myh4. We further observed that cachexia-related gene activation was restricted to myonuclear subsets and did not overlap with the downregulation of differentiation-related genes. In conclusion, our analysis uncovered early disruption of myofiber transcriptional identity in mice, characterized by the selective loss of *Maf* and *Myh4,* alongside myonuclear-specific activation of cachexia-related genes; a conserved feature in both murine and human PDAC cachexia.

## Results

### snRNAseq reveals PDAC disrupts the skeletal muscle microenvironment prior to overt muscle wasting

First, we defined our murine model of cancer cachexia. Tissues were collected every three days starting at day 6 (Figure 1A). PDAC tumors were detectable by day(D) 12 (Figure.1B). Body weight decreased (5.9%) starting at D12 compared to sham (Figure 1C). At our last time point, body weight decreased by 15.5%. PDAC tumors induced quadriceps muscle loss at D15 and D17/18; −14% and −23% respectively (Figure 1D and Supplemental Figure 1). Taken together, these data validate our murine model of cancer cachexia. Next, we defined the muscle microenvironment prior to muscle wasting. We performed snRNAseq on the D12 quadriceps muscle. First, we performed unbiased clustering that identified 13 clusters (Supplemental Figure 1). The same cell types were combined to identify 9 unique cell types, with the predominant cell type being myonuclei and fibro-adipogenic progenitor cells (FAPs) (Figure 1E,F). Cell types were confirmed by known cell type gene markers (Figure 1G). To determine if PDAC disrupts the muscle’s microenvironment, we compared the proportion of cell types between sham and PDAC and identified increased myonuclei, reduced FAP and macrophage proportions without changes in the satellite cell proportions. These data provide evidence that PDAC tumors disrupt the skeletal muscle microenvironment prior to overt muscle wasting.

**Figure 1:**
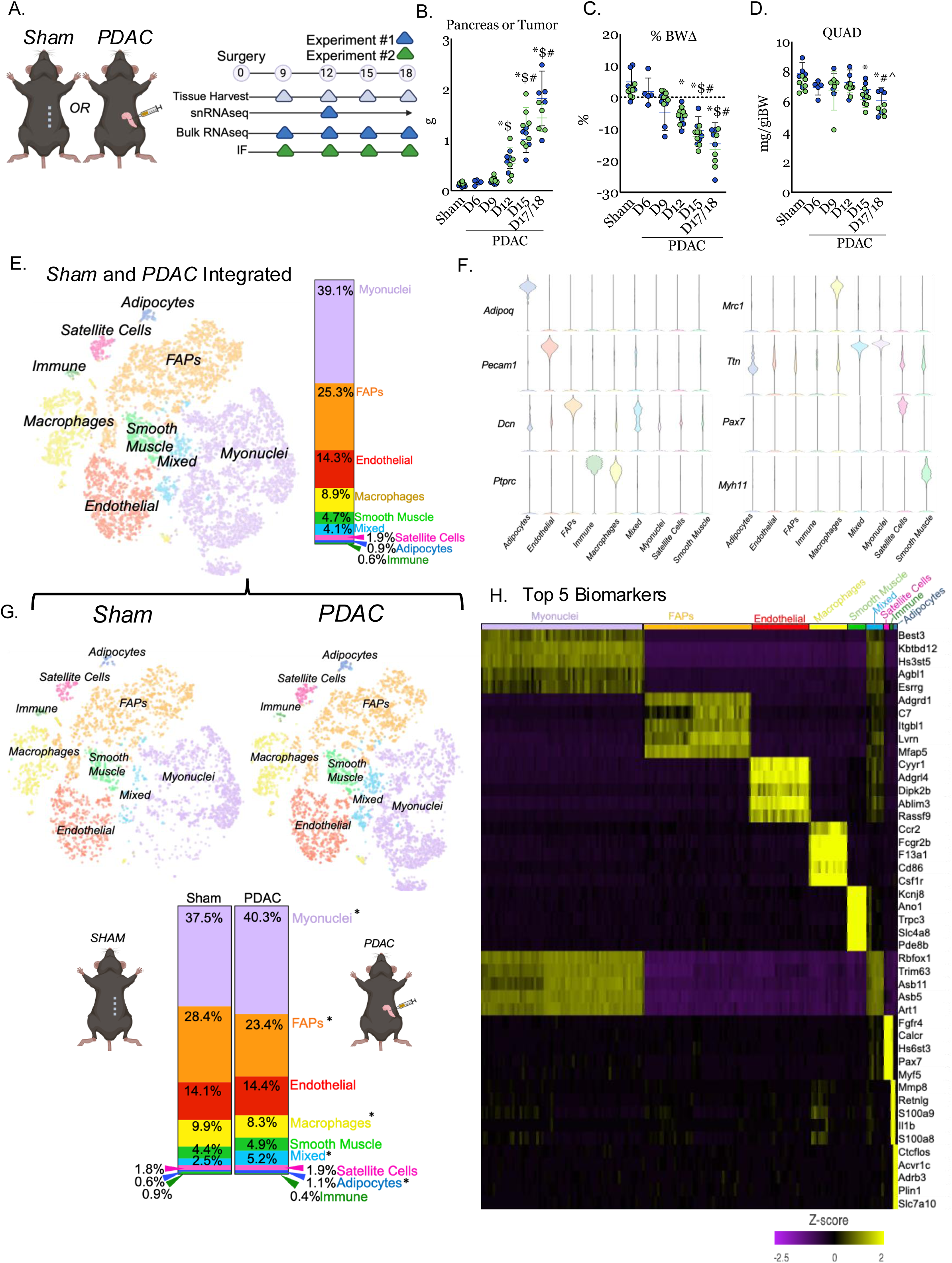
snRNAseq reveals PDAC disrupts the skeletal muscle microenvironment prior to overt muscle wasting. **A)** Study design. Two separate time course experiments were completed to provide sufficient tissue for single nucleus and bulk RNAseq and immunofluorescence. 10-week-old male C57BL/6J mice were orthotopically injected with 50,000 KPC-32908 cells into the pancreas (PDAC) or underwent a sham surgery (sham). Mice were euthanized every 3 days starting on D6 until D18. **B)** Pancreas or tumor mass across the time course. Tumors were observed starting D12. **C)** Body weight change (%BWΔ) at the time of euthanasia accounting for tumor mass. **D)** Quadriceps (QUAD) mass relative to initial body weight (IBW). Kruskal-Wallis with Dunn’s multiple comparison tests (**B**) and one-way ANOVA using Tukey’s multiple comparison tests (**C**, **D**) were used to compare tissue and body weight across the time course. *Different from sham, $different from D6, #different from D9, ^different from D12. P<0.05. **E)** T-SNE plot of integrated sham and PDAC skeletal muscle cells from snRNAseq of the quadriceps. Proportion of cell types in the integrated sham and PDAC samples. **F)** Violin plot of genes used to identify cell types. Cell type biomarkers are listed in Tables 1 and 2 and Supplemental Figure 2. **G)** T-SNE plot from **E** separated by condition. Proportion of cell types in the sham and PDAC. *P<0.05. **H)** Heat map of Z-scores for the top 5 biomarkers in each cell type.

### snRNAseq reveals PDAC induces heterogeneous expression of myosin heavy chains prior to muscle wasting

Next, we interrogated the skeletal muscle myonuclei population in PDAC. First, we investigated skeletal muscle structural genes, as prior studies showed that cancer cachexia reduces skeletal muscle structural proteins[21, 22]. Our snRNAseq revealed that at the pseudo-bulk (all cell types) and myonuclei level, *Myh4* was decreased while *Myh1* and *Myh2* were increased in PDAC vs sham (Figure 2A). Mature muscle differentiation transcription factors, *Ckm* and *Maf*, were also reduced in PDAC vs sham. Next, we interrogated genes enriched in fiber types identified IIA, IIX, or IIB[23] in our myonuclei population. Our snRNAseq reveals that genes enriched (red up arrow) in fiber types IIA and IIX are increased in PDAC vs sham. Genes enriched in type IIB fibers are decreased in PDAC vs sham (Figure 2B). Altogether, these data suggest a loss of type IIB fiber transcripts and increased fiber type IIA and IIX associated transcripts prior to wasting.

**Figure 2:**
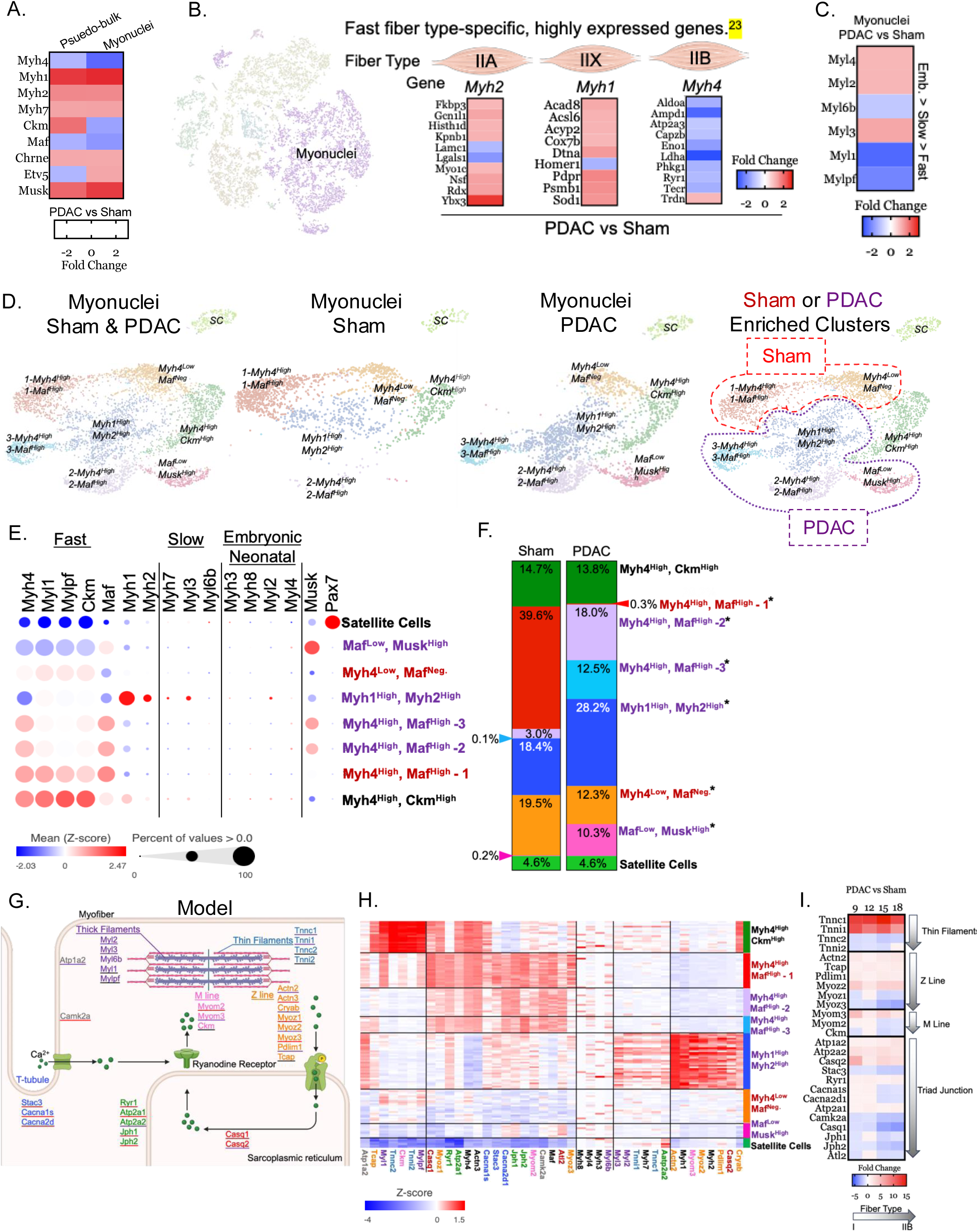
snRNAseq reveals PDAC induces heterogeneous expression of myosin heavy chains prior to muscle wasting. **A)** Heat map of myosin heavy chain and mature skeletal muscle genes in the snRNAseq data by pseudo-bulk (all cell types) and myonuclei comparing PDAC vs sham. Data are expressed as fold change. **B)** Heat map of fast fiber type-specific expressed genes in the myonuclei sub-population comparing PDAC vs sham. Data are expressed as fold change**. C)** Heat map of myonuclei myosin light chain genes comparing PDAC vs sham. Genes are listed in order of developmental stage, from embryonic to fast fiber type. **D)** T-SNE plots of myonuclei sub-populations in the integrated sham and PDAC, each condition, and the integrated plot highlighting the populations enriched in each condition. Red identifies sham-enriched, and purple identifies PDAC-enriched myonuclei sub-populations. **E)** Skeletal muscle fiber type gene expression and percentage of nuclei expressing the genes. The high or low expression of the myosin heavy chains, *Maf*, and *Ckm* was used to label myonuclei sub-populations. Z-scores are expressed by bubble color. **F)** Proportion of myonuclei sub-populations in the sham and PDAC. *Proportion of cell type sin the myonuclei sub-population difference between sham and PDAC. P<0.05. **G)** Modified image from biorender illustrating the components of striated skeletal muscle necessary for function. **H)** Hierarchical clustering of myonuclei sub-populations by striated skeletal muscle genes. I) Heat map of striated skeletal muscle structural genes across the time course. Genes are sorted from top (white) to bottom (grey), having greater expression in fiber type I to IIb fibers. Data are expressed as fold change.

We performed unbiased clustering of the myonuclei population to identify 7 myonuclei sub-populations (Figure 2D). Sup-populations were named by established muscle associated genes: myosin, musk, and Pax7 gene expression (Figure 2E). Notably, there are two populations enriched in sham (red outline and text) and 3 sub-populations enriched in PDAC (purple outline and text) with one sub-population only in PDAC (Figure 2D). Our data suggest that sham enriched sub-population *Myh4^High^,Maf^High^-1* expands to make PDAC *Myh4^High^,Maf^High^-2 and Myh4^High^,Maf^High^3,* because of the gradual reduction in Myh4, (Figure 2E), hierarchical clustering of maturation genes in these sub-populations (Supplemental Figure 2B), and trajectory analysis (Supplemental Figure 4B).To understand the sub-population expansion in PDAC, we next sought to define if the skeletal muscle contractile transcriptome can explain these sub-population differences (Figure 2G). We identify that sham *Myh4^High^,Maf^High^-1* expansion into PDAC *Myh4^High^,Maf^High^-2 then Myh4^High^,Maf^High^-3, then Myh1^High^, Myh2f^High^* is partially explained by a loss of mature muscle transcriptome markers (Figure 2H). At the bulk RNAseq level, PDAC cachexia reduced type IIB fiber transcription factors that regulate skeletal muscle function and structure (Figure 2I). These data suggest that healthy muscle (sham) expresses numerous myonuclei sub-populations that become more heterogeneous in PDAC.

### Muscle differentiation identity factor Maf and its target Myh4 are lost while Myh1 and Myh2 are gained during the progression of muscle wasting in pancreatic cancer cachexia

Our prior figures revealed *Myh4* gene expression loss in PDAC. First, we defined how the transcription factors enriched in fully differentiated fibers [8] are reduced during cachexia progression (Figure 3A and Supplemental Figure 2E). Recent work identified the direct link between *Myh4* and *Maf [8, 24]*; therefore, we sought to determine if we observed a similar link between in PDAC cachexia. First, we identify a loss of *Maf* gene expression during the timecourse of PDAC cachexia (Figure 3B). This loss in *Maf* gene expression is specific to large *Maf/Mafc* and *Mafa* (Supplemental Figure 2F). The loss of *Maf* gene expression was positively correlated with greater body weight loss (Figure 3C). We interrogated myosin heavy and light chains and *Maf* gene expression in PDAC vs. sham across the cachexia time course and reported a reduction in *Myh4*, *Myl1*, and *Mylpf* gene expression (Supplemental Figure 2G). The reduction in *Maf* is positively correlated to the reduction in *Myh4* gene expression (Figure 3D). Next, we interrogated the protein expression of MYH4 and MYH2 by immunofluorescence and gene expression of *Maf* by RNAscope in skeletal muscle longitudinal sections. Our data identifies that fibers expressing MYH4 overlap with *Maf* puncta (Figure 3F). Additionally, we observe an increase in MYH1 and 2 expressing fibers which occurs longitudinally across the entire fiber (Figure 3G). Next, we identified the gene expression of known Maf targets in our snRNA and bulk RNAseq data (Supplemental Figure 2F). We show a positive association of Maf targets to mature muscle markers (Figure 2E and Supplemental Figure 2G), further supporting that the relationship between Myh4 and Maf is maintained cachexia. Altogether, these data suggest that PDAC cachexia drives the loss of Maf and downstream target Myh4 while simultaneously gaining Myh1 and Myh2 expression.

**Figure 3:**
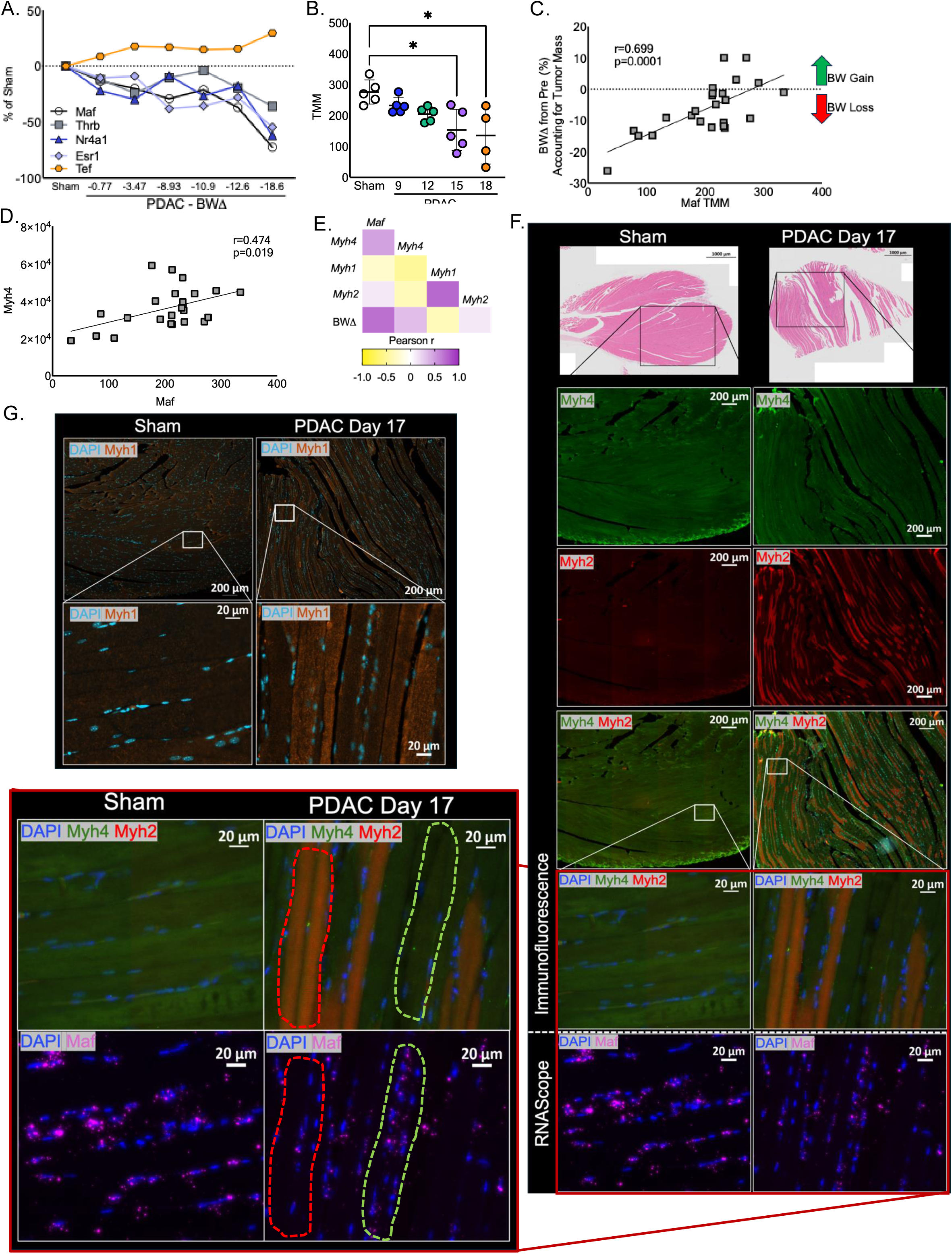
Muscle differentiation identity factor Maf and its target Myh4 are lost while Myh1 and Myh2 are gained during the progression of muscle wasting in pancreatic cancer cachexia. **A)** Transcription factors enriched in fully differentiated fibers from Dos Santos et al. [8]. **B)** *Maf* gene expression during the progression of cachexia. Data are expressed as trimmed mean of M values (TMM). One-way ANOVA was used to compare normalized counts. *P<0.05. **C)** Association of body weight change from pre, accounting for tumor mass, associated with *Maf* TMM counts. Statistical test: Pearson correlation coefficient. **D)** Association of *Myh4* TMM gene counts with *Maf* TMM gene counts. Statistical test: Pearson correlation coefficient. **E)** Heat map of body weight change from pre associated with myosin heavy chains. Statistical test: Pearson correlation coefficient. **F)** We performed immunofluorescence (IF) to determine protein expression of MYH4 and MYH2 and RNAscope to determine gene expression of *Maf* in the sham and PDAC D17 condition. Images were enlarged to highlight the overlap between MYH4 and *Maf*. In the enlarged image of PDAC D17, we outlined the areas having high MYH2 and low *Maf* expression in red and high Myh4 and *Maf* expression in green. **G)** Immunofluorescence of Myh1 in the sham and PDAC D17 condition. Immunofluorescence was completed in longitudinal sections of the gastrocnemius. PDAC D9, 12, and 15 can be found in Supplemental Figure 3. We first performed Myh7 expression to identify fibers not expressing Myh7 to prevent bias by imaging the more oxidative portion of the gastrocnemius muscle.

### PDAC alters oxidative phenotypes in a subgroup-specific fashion with a lack of evidence to support a fiber type shift in the myonuclei by snRNAseq or in muscle by bulk RNAseq

Given the similar mature myosin heavy chain expression, we sought to further define the myonuclei sub-populations. As expected, sham-enriched sub-populations are enriched in GO biological process of glycogen metabolism, TCA cycle, myotube cell development, and actin-mediated cell contraction (Supplemental Figure 4A). In the PDAC sub-populations, we observe GO biological process pathways of fatty acid beta-oxidation, steroid hormone-mediated signaling, and circadian behavior; to name a few (Supplemental Figure 4A). The PDAC only sub-population is enriched in selective autophagy, IL-6-mediated signaling, and negative regulation of stem cell differentiation pathways. Based on the transition of myosin heavy chains in the prior figures, we speculated that we would observe a gradual expansion of oxidative genes and a reduction in glycolytic genes. In contrast, we observed the greatest induction of oxidative genes in the PDAC enriched *Myh1^High^,Myh2^High^*population (Supplemental Figure 4C). We observe the greatest induction of glycolytic genes in the *Myh4^High^,Ckmf^High^* sub-population which is equally proportional in PDAC and sham (Supplemental Figure 4D). These findings agree with our bulk RNAseq data showing a lack of transition to an oxidative (Supplemental Figure 4F) or glycolytic (Supplemental Figure 4G) gene signature during PDAC cachexia progression. Furthermore, we do not observe a sub-population transition from glycolytic to oxidative in our snRNA and bulk RNAseq (Supplemental Figure 4E,H). Our data also reveals low evidence for PDAC effects on skeletal muscle regeneration, fibrosis, or proliferation (Supplemental Figure 5). In summary, these data identify that PDAC alters oxidative phenotypes in a sub-population-specific manner instead of a homogeneous transition across all populations.

### Induction of the cachexia gene program is heterogeneous and distinct from myosin isoform identity

Next, we sought to determine if the molecular cachexia gene signature drives the myosin heavy chain isoform expression. First, we culminated 22 skeletal muscle cachexia studies and defined the top cachexia genes (Tables 4 and 5). By bulk RNAseq, we observe a gradual increase in most of the top 30 cachexia genes during the progression of PDAC cachexia (Figure 5A). At single-nucleus resolution, most of the genes are upregulated at the pseudo-bulk and myonuclei level (Figure 5B). Most surprisingly is that the PDAC only sub-population - *Maf^Low^,Musk^High^* – robustly upregulates most of the cachexia genes when compared to all the myonuclei sub-populations (Figure 5C). These data illustrate a new cachexia myonuclei population in PDAC. First, we illustrate the expression of *Myh 4, 2, 1*, and *7* in PDAC only myonuclei sub-populations (Figure 5E) and a few selected cachexia genes (Figure 5G). We then set out to illustrate if the cachexia gene signature overlapped with mature muscle fiber gene markers. We illustrate that the cachexia gene signature does not uniformly overlap with mature differentiated fiber genes (Figure 5F). We performed gene rank sum of differentiation and cachexia genes. PDAC cachexia decreased the sum differentiation gene rank and increased cachexia gene rank (Figure 4H). At this pre-cachexia phase, there is a modest association between reduced differentiation genes and increased cachexia genes (Supplemental Figure 2I). In summary, we show that the cachexia gene signature is robustly upregulated prior to overt wasting, is ubiquitously expressed, and the shift in myosin expression modestly overlaps with cachexia genes (Figure 5H).

**Figure 4:**
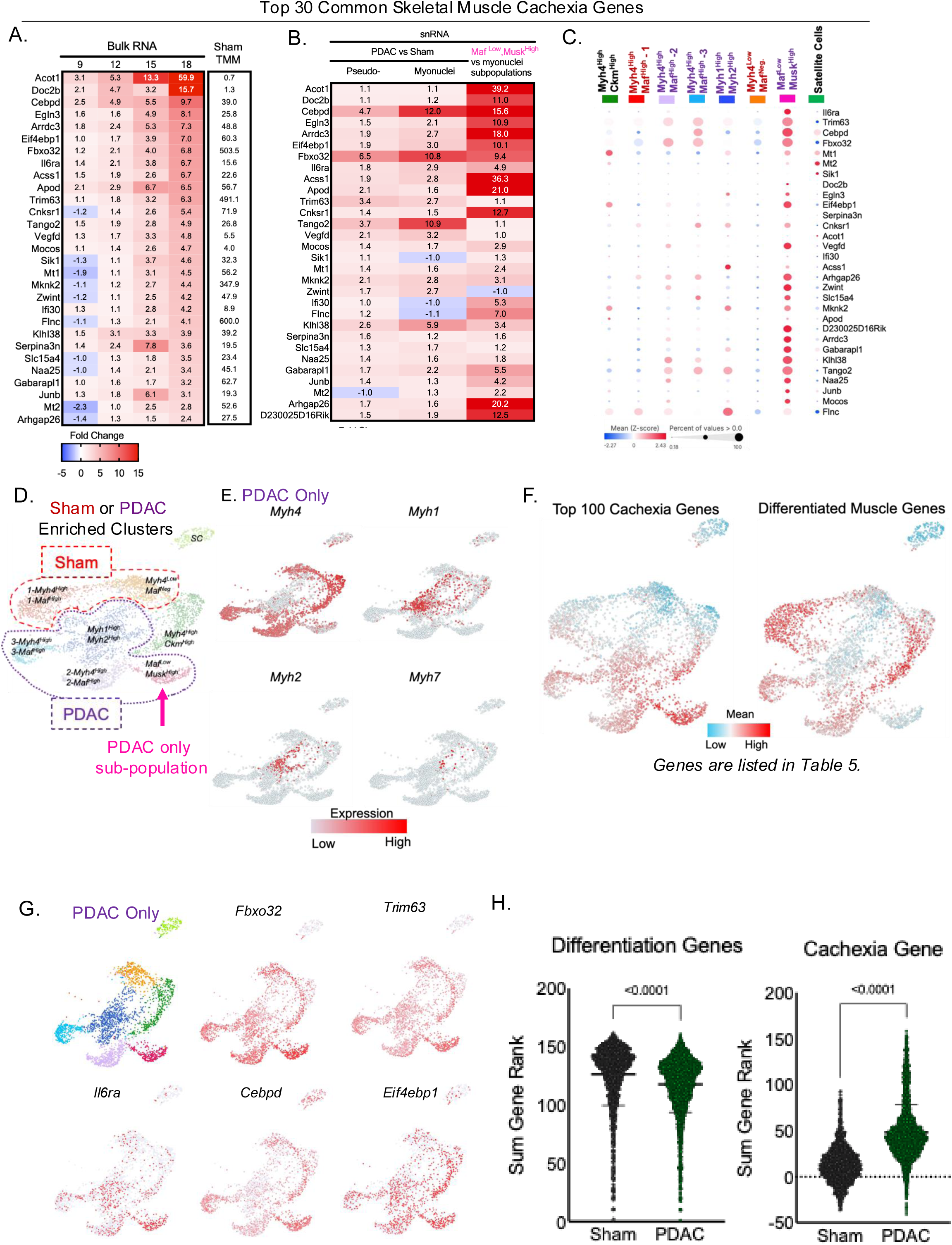
Induction of the cachexia gene program is heterogeneous and occurs distinctly from myosin isoform identity. **A)** Heat map of cachexia genes in bulk RNAseq across the time course comparing PDAC vs sham. Data is expressed as fold change. **B)** Heat map of cachexia genes in the snRNAseq data by pseudo-bulk (all cell types) comparing PDAC vs sham, myonuclei comparing PDAC vs sham, and myonuclei PDAC only sub-population *Maf^Low^,Musk^High^* vs all myonuclei sub-populations. Data are expressed as fold change. **C)** Gene expression and percentage of nuclei expressing top upregulated cachexia genes in snRNAseq. **(A-C)** Cachexia genes were identified using BaseSpace Illumina to curate the top upregulated genes across several models of murine cancer cachexia (Tables 4 and 5). **D)** T-SNE plot of myonuclei sub-populations in the integrated sham and PDAC highlighting the populations enriched in each condition. Red identifies sham-enriched, and purple identifies PDAC-enriched myonuclei sub-populations. The pink arrow identifies the PDAC only sub-population. **E)** T-SNE plots of *Myh 4, 1, 2, and 7* in myonuclei sub-populations in PDAC. **F)** T-SNE plots of the top 100 cachexia genes (Tables 4 and 5) and differentiated gene markers in the integrated sham and PDAC, identifying the sub-populations of highest expressing cachexia genes and lack of overlay with differentiation genes. **G)** T-SNE plots of *Fbxo32, Trim63, Il6ra, Cebpd, and Eif4ebp1* in myonuclei sub-populations in PDAC. **H)** Sum gene rank of differentiation and cachexia genes in Sham and PDAC conditions from single nucleus RNAseq.

**Figure 5:**
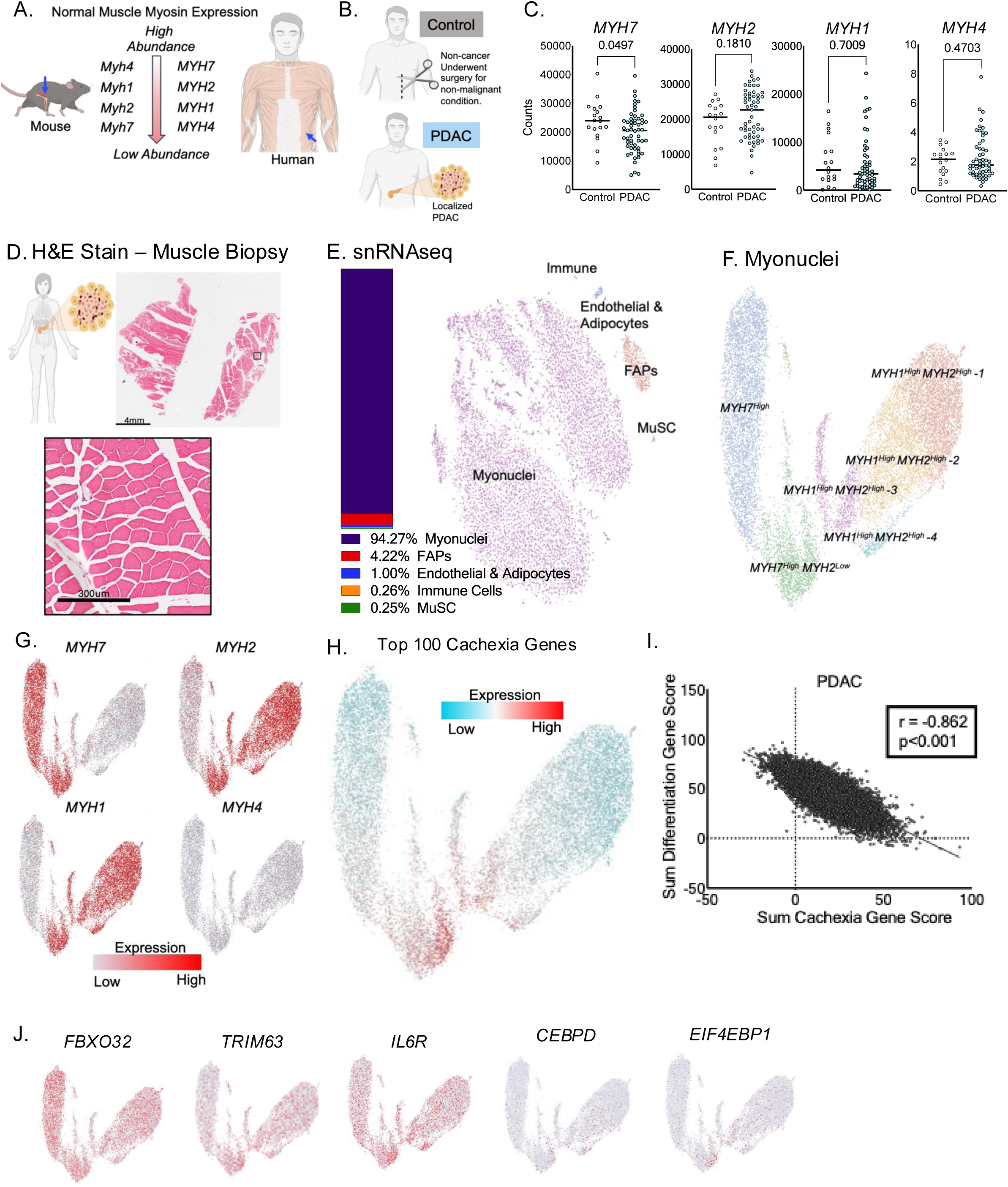
Patients with pancreatic cancer show myosin dysregulation and discrete activation of cachexia genes in skeletal muscle. **A)** Illustration of mature myosin heavy chain abundance in mice and human skeletal muscle. **B)** Illustration of treatment groups for the human muscle biopsy samples. Rectus abdominus muscles were taken from patients in the control condition who did not have cancer but underwent surgery for a non-malignant condition. PDAC patients had localized cancer and underwent surgery to remove the tumor at the time of biopsy. **C)** Myosin heavy chain isoform gene expression from bulk RNAseq. Statistical test: Unpaired t-test. **D)** H&E section from the PDAC patient muscle biopsy. The muscle sample was used for snRNAseq. **E)** UMAP plot of muscle cell types from snRNAseq in one patient with PDAC cancer cachexia. **F)** UMAP of myonuclei sub-populations from snRNAseq. **G)** UMAPs of myonuclei sub-population myosin heavy chain gene expression. H) UMAP of the top 100 cachexia genes. **I)** Association of the sum differentiation gene rank with cachexia gene rank in single nucleus RNAseq. **J)** UMAPS of *FBXO32*, *TRIM63*, *IL6RA*, *CEBPD*, and *EIF4EBP1* in myonuclei sub-populations.

### Patients with pancreatic cancer show myosin dysregulation and discrete activation of cachexia genes in skeletal muscle

We sought to determine if the heterogeneous cachexia gene program and myosin isoform identified in our pre-clinical model translated to pancreatic cancer patients. First, we illustrate the differences in myosin heavy chain isoform abundance in mice vs humans (Figure 6A). Muscle biopsies from the rectus abdominis were taken from patients undergoing surgery for non-malignant conditions (control) or cancer patients undergoing surgery for localized PDAC (Figure 6B). We performed bulk RNAseq on the muscle biopsy from control and PDAC patients (Figure 6C). PDAC patients have decreased *MYH7* expression compared to controls, illustrating a reduction in mature myosin isoform. Next, we performed snRNAseq on one female patient with the highest body weight loss (−31%) in our patient cohort and where the biopsy contained predominately skeletal muscle (Figure 6D). We had sufficient nuclei from all cell types identified (Supplemental Figure 6E), ∼94% of the nuclei captured were myonuclei (Figure 6E), and the myonuclei sub-populations segregate by myosin heavy chain isoform expression (Figure 6F,G). When we overlap the top 100 cachexia genes, we observe heterogeneous expression of cachexia genes that are not specific to one myosin isoform (Figure 6H). Next, we interrogated a few cachexia genes to further illustrate the heterogeneity (Figure 6J) supporting our prior observation in our pre-clinical model of PDAC cachexia, showing myonuclei myosin dysregulation and heterogeneous activation of cachexia genes in pancreatic cancer cachexia. We performed gene rank sum of differentiation and cachexia genes, and show that at this late cachexia stage, with severe muscle wasting there is a strong negative association between differentiation and cachexia gene (Figure 5I). In summary our data suggest that the fate of myonuclei occurs early in cachexia such that (1) myonuclei remain unaffected by the cancer environment until late-stage cachexia, (2) myonuclei express cachexia gene signature or a dedifferentiation gene signature, or (3) myonuclei exhibit both a cachectic and dedifferentiation gene signature prompting irreversible changes (graphical abstract).

## Materials and Methods

### Mice

We purchased fifty-five 10-week-old male C57Bl/6J mice from Jackson Lab. We completed two independent experiments: (1) at Indiana University and (2) at Oregon Health and Science University. There were no differences in animal characteristics and cachexia development between universities. Mice were group-housed in a conventional housing facility with ad libitum access to autoclaved food (Envigo) and sterile water and maintained on a 12hr light/dark cycle. Mice were acclimated to the facilities for two weeks prior to being randomized into experimental groups. Mice were euthanized every three days, starting on D6. Sham mice were euthanized on day 9 (Exp.#1) or 18 (Exp.#2). To control for cage conditions, mice were group housed with each experimental timepoint allocated to each cage. Mice did not reach protocol approved euthanasia requirements of <5% body fat or > 20% body weight loss. Orthotopic pancreatic tumors of KPC32908 cells rarely have an excessive tumor burden prior to reaching our approved euthanasia requirements and were not reached in our animal studies. These studies were performed in accordance with Indiana University School of Medicine and Oregon Health and Science University’s Institutional Animal Care and Use Committees and approved in protocol numbers 19036 (IU) and 4677 (OHSU).

### Orthotopic Implantation of KPC Cells

Extensive surgical procedures for orthotopic implantation of pancreatic cancer cells have been described elsewhere[18]. Briefly, 12-wk-old male mice were anesthetized under isoflurane and subcutaneously injected with Ethiqua Extended-Release 3.25mg/kg immediately before surgery (Exp. #1), or 5mg/kg meloxicam in sterile PBC before surgery and then at 24hrs from the first injection (Exp. #2). The pancreas was injected with 5×10^4^ KPC32908 in 40µl of sterile PBS for 20secs. The spleen and accompanying pancreas were gently laid back into the abdominal cavity using ring-tip forceps. The muscle wall was closed using sutures, and the skin flap was closed with sterile wound clips. Sham mice underwent the same procedures without manipulation of the spleen or pancreas. Analgesics used followed individual institutional animal care requirements. There were no differences in endpoint animal characteristics and cachexia development between experiments.

### IRB

The study was conducted under the Indiana University Institutional Review Board (IRB) approved protocol #1312105608, the same protocol referenced in our prior work[25, 26].

### Cell Culture

KPC 32908 tumor cells (provided by Dr. David Tuveson; Cold Spring Harbor, NY), were isolated from a pancreatic tumor in a male LSL-KrasG12D:LSL-Trp53R172H:Pdx1-Cre (KPC) mouse on a C57BL/6 background[27]. KPC cells between passages 4-6 were thawed at 37°C in a water bath and grown using DMEM growth medium supplemented with 10% FBS and 1% pen-strep. Cells were seeded at 5X10^5^ and allowed to grow for 48 hours. In preparation for surgery, the cells were detached from the plate using TripLE reagent and were counted using a Countess. Cells were spun at room temperature at 1500rpm for 5mins and an adequate number of cells were resuspended in sterile PBS for tumor cell injections. Mice injected with KPC32908 tumors are referred to as PDAC throughout the manuscript.

### Single-Nuclei Isolation and Sequencing – Mouse Muscle

#### Nuclei Isolation

We isolated nuclei from the quadriceps muscle of 4 sham mice and 4 PDAC D12 mice. Approximately 50mg of flash-frozen quadriceps muscle tissue was powdered in liquid nitrogen using a mortar and pestle. Samples were then dounced on ice in 7ml of NPB-VRC (0.25M sucrose, 0.01M Hepes pH7.5, 0.01M KCL, 0.1%NP40, 0.01M VRC added at RT, 1X halt protease inhibitor, 0.001M DTT and 1X RNase inhibitor) 10 times. Samples were spun at 200xg at 4°C for 10mins in a swinging bucket centrifuge. The supernatant was removed and 1ml of PBS-N-VRC was added to gently disturb the pellet. Samples underwent a second spin of 200xg at 4°C for 5mins and were resuspended in 1000µl of PBS-N-VRC, then strained through a 40µm filter. Nuclei were FACS sorted using DAPI, and we collected ∼500,000 nuclei/sample. After sorting, samples were suspended in 1000µl of PBS-N-VRC and spun 200xg for 10 minutes. Samples were resuspended in 30µl of PBS-N. Samples were counted at the IU School of Medicine Genomics Core; the sham and PDAC D12 samples were resuspended, pooled, and submitted for sequencing.

#### Sequencing

Single-nuclei 3’ RNA-seq assay was conducted using the 10x Chromium single-cell system (10x Genomics, Inc). Each single-nuclei suspension was first checked for nuclei quality and quantity. The four samples were pooled for each condition. Single-cell gel beads in an emulsion containing barcoded oligonucleotides and reverse transcriptase reagents were generated with the Next Gem single-cell reagent kit (10X Genomics). cDNA was synthesized and amplified. The quality of cDNA and library were examined by Bioanalyzer at each step. The final dual-indexed library was sequenced on an Illumina NovaSeq 6000. 100-bp reads, including cell barcode and UMI sequences were generated.

#### Filtering and analysis

10X chromium fastq files were uploaded to Partek Flow. First, we filtered by excluding nuclei with less than 200 or more than 5000 genes, and nuclei with more than 5% of mitochondria (Supplemental Figure 1C,D). Then, counts were normalized to counts per million, and genes with a value of 0 in 100% of the nuclei were removed. See supplemental text for Partek command lines (Supplemental Figure 8). Extended methods can be found in Supplemental Methods.

### Bulk RNA Isolation and Sequencing

#### RNA isolation

RNA was isolated from ∼50mg of snap-frozen quadriceps tissue using the miRNeasy Mini Kit (Qiagen, cat: 217004). The concentration and quality of total RNA samples were assessed using Agilent 2100 Bioanalyzer, and an RNA Integrity Number (RIN) of eight or higher was required to pass the quality control. We isolated RNA from 4-5 mice/time points. Library preparation and sequencing were completed at the Massively Parallel Sequencing Shared Resource at Oregon Health and Science University using the Illumina NovoSeq6000. 150ng of total RNA was used for each sample. RNA sequencing cDNA library preparation, sequencing, and filtering details are listed in the extended methods.

### Histology

#### Tissue Preparation

Tibialis anterior and gastrocnemius muscles were excised from the hindlimbs and placed in 1.5ml tubes with 1ml of 10% neutral buffered formalin. Tubes were placed on a rocker at 4℃ for 48 hours. Samples were washed twice with cold PBS and placed in a 1.5ml tube of 1ml 70% ETOH at 4℃ until ready for processing. Tibialis anterior muscle samples were cross-sectioned, and gastrocnemius samples were embedded longitudinally; all were sectioned to a thickness at 5um. Samples were paraffin processed on the Tissue Tek VIP (vacuum infiltration processor. Samples were embedded and stained for H&E and Trichrome at the Histology Shared Resource at Oregon Health and Science University.

#### Immunohistochemistry and Immunofluorescence

Tibialis anterior muscle samples were stained for H&E and Trichrome. The Leica Autostainer XL was used for automated H&E staining using Hematoxylin 7211 and Alcoholic Eosin Y. Immunofluorescence staining was performed on gastrocnemius longitudinal muscle sections. Paraffin-embedded slides were baked at 65°C for 15 minutes, then slides were deparaffinized. Next, we performed antigen retrieval via instant pot with Tris-EDTA pH9 buffer for 10 minutes. Slides were incubated in 0.1M glycine, and 0.1% Triton-X for 10 minutes. Slides were washed with PBS and then treated with TrueBlack Lipofuscin autofluorescence quencher for 30 seconds. Slides were washed, then blocked with blocking buffer: 1:25 dilution of M.O.M (Vector Labs, cat: MKB-2213-1) and 5% goat serum in PBS, at RT for 1hr. Next, slides were incubated overnight at 4°C in the dark with primary antibodies. Primary antibodies were diluted 1:100 for Myh4, Myh2, and Myh1 (Developmental Studies Hybridoma Bank, cat: BF-F3, SC-71, and 6H1-S). Slides were washed, then incubated with secondary antibodies for 1hr at RT. Secondary antibodies were diluted 1:400 Alexa Fluor 488:IgM, 647:IgM, or 555:IgG1. Slides were washed with PBS and incubated with 1ug/ml DAPI for 5 minutes. Slides were washed, then mounted with Prolong Gold Antifade and stored at 4°C in the dark until imaging.

#### RNAscope

In-situ hybridization was performed for Maf using the Advanced Cell Diagnostics (ACD) multiplex fluorescent staining kit (cat: 322800) on the Leica Bond Rx automated stainer. Slides were deparaffinized in two five-minute incubations in xylenes, placed in two three-minute incubations of 100% ethanol to remove the xylenes, and rehydrated in RNase-free ddH2O. To reduce auto-fluorescence, slides were incubated in a high pH hydrogen peroxide solution for 45 minutes while being exposed to bright light. Slides were then loaded onto the Bond for onboard pretreatment of epitope retrieval (15mins at 95 °C with Lecia Bond ER2 (cat: AR9640)) and a mild protease treatment (ACD Bio Protease III diluted 1:20 in RNase-free PBS for 15mins). A solution of Maf was dispensed and incubated for 2hrs at 60°C, followed by manufacturer instructions for amplification steps. Slides were then unloaded, incubated with Biotium Tyramide CF568 diluted 1:500 in TSA Amplification Diluent, and then loaded back onto the Bond for an HRP block and subsequent amplification steps for channel 2 of the kit. Once unloaded, Biotium Tyramide CF640 diluted 1:500 in TSA Amplification Diluent was applied for 8mins, followed by nuclear detection with DAPI 1:10,000 in ddH2O. Slides were cover slipped using Prolong Gold.

### Microscopy

#### Slide scanning

Sample images were acquired on a Zeiss Axioscan 7 slide scanner. Fluorescence images were illuminated with a Colibri7 LED light source (488nm, 555nm, 647nm) and captured on a Hamamatsu Orca Flash 4.0 camera (0.324um/pixel) using a 20x 0.8NA objective. Brightfield images were obtained with a 10x 0.45NA objective on a Zeiss Axiocam 705 color camera (0.346um/pixel).

### Single-Nuclei Isolation and Sequencing – Human Muscle

#### Nuclei Isolation

Briefly, 50mg flash-frozen muscle stored at −80°C was used for source material. 750µl TsT nuclei lysis buffer was added to 50mg of flash frozen tissue in a 1.7ml tube. The minced tissue was transferred to a pre-chilled dounce homogenization tube. Following several rounds spins and straining, the supernatant was removed, and nuclei were resuspended in 200µl NSB. Nuclei quality was assessed visually and quantitatively using trypan blue, Hoescht, and the Invitrogen EVOs scope and Countess II FL. Full isolation methods can be found in the Extended Methods. Fresh nuclei were resuspended (3400/µl) in NSB and used immediately for library preparation using the Fluent PIPseq V T10 3’ Single Cell RNA Kit according to the manufacturer’s protocol. Pre-made library and partial lane sequencing was performed by Novogene using the NovaSeq X Plus Series PE150, Q30>85%. Raw FASTQ files were pre-processed with PIPseeker and then aligned to the human reference genome GRCh37/hg19 with STAR. Aligned reads were traced to individual cells and gene expression levels were quantified based on the number of UMIs detected. The filtered barcode matrices were used for further analysis. Cells were retained if they had less than <30,000 total reads. Here, due to the nature of single nucleus data, we don’t expect mitochondria reads. Doublets were removed with scDblFinder[28, 29].

#### Filtering and analysis

Filtered CSV files were uploaded in Partek Flow. We performed Nuclei with >7000 genes were excluded, and then we filtered out nuclei with ≦ 0 genes in 99% of nuclei for downstream analysis (Supplemental Figure 6D). Then, counts were normalized to counts per million. See supplemental text for Partek command lines (Supplemental Figure 8). We performed a PCA using the top 2000 features, then computed graph-based clusters and biomarkers on 100PC as we accounted for >99% of the variance. Cell types were identified based on the expression of validated cell-type genes (Supplemental Figure 6E).

### Statistics

One-way ANOVA with Tukey’s post hoc test or Kruskal-Wallis test with Dunn’s multiple comparison post hoc test was performed when necessary; as noted in the figure legends. One-way ANOVAs were used for comparison of the means of tissue weights and RNAseq normalized counts. Unpaired two-tailed Student’s tests were used for the comparison of gene expression from human cancer and control patients. Cell type proportion analysis was determined by calculating the ratio of a cluster’s count by the total nuclei count within the same condition. Two-sided statistical tests were used to assess differences in these proportions between conditions. When both cluster counts were ≥30 in each condition, a two-proportion Z-test with pooled variance was applied. When either count was <30, Fisher’s exact test was used instead to avoid normality assumptions. To control the false discovery rate across clusters within each sub-population, p-values were adjusted using the Benjamini-Hochberg procedure. Statistical significance was defined as an adjusted p-value < 0.05. All analyses were conducted in Python (version 3.11.5) using SciPy (version 1.11.3). The differences between group means were considered statistically different when P<0.05. Data are presented as mean ± standard deviation. All statistical analyses were performed using GraphPad Prism version 10.0.

## Discussion

Our previous understanding of the transcriptional and proteomic changes in skeletal muscle during cancer cachexia has largely been based on whole-tissue analyses. Recent snRNAseq studies in cachectic mice have identified distinct catabolic myonuclei populations[19, 20]. However, these analyses were conducted in wasted muscle. To address this knowledge gap, we examined transcriptional changes at the single-nucleus resolution prior to overt muscle wasting. We observe a progressive loss of muscle differentiation identify factor *Maf* and its target *Myh4*, accompanied by increased expression of *Myh1* and *Myh2* during cachexia progression. Notably, while we identify a cachexia-only myonuclear subset, the expression of cachexia genes occurred heterogeneously across these sub-populations. Importantly, the loss of muscle identity and the gain of catabolic gene expression occurred in distinct myonuclear populations, a pattern that was also present in skeletal muscle in patients with PDAC cachexia.

Adult mouse hindlimb skeletal muscles predominately express *Myh4*, with much lower levels of *Myh1* and *Myh2*, which are considered less mature fast fibers[30, 31]. Recent studies have identified *Maf* as part of the differentiated fiber gene repertoire [8]. Large Maf proteins, including c-Maf, promote the expansion of type IIb fibers by binding to MYH promoters and driving Myh4 expression[24]. In our analysis, *Maf*/c-Maf exhibited the highest RNA abundance among the large Mafs but was significantly reduced during cachexia, suggesting that *Myh4* downregulation is likely due to diminished *Maf* transcriptional activity. Prior work has shown that in myofibers, calcium signaling can induce *Maf* expression[8], while IL-6 stimulation promotes *Maf* expression in T cells[32], highlighting its broader role in differentiation processes. Although it is tempting to speculate that overexpressing *Maf* could restore *Myh4* expression and support muscle fiber identity, the pleiotropic roles of *Maf* in multiple cell types warrant caution in pursuing such an approach.

Total MYH protein expression is reduced in multiple murine models of cancer cachexia (C26, LLC, PDAC)[21, 22, 33, 34]. While fiber size changes are accompanied by a shift in myosin isoform expression[35, 36], findings in cachexia are mixed: some report changes[37], others do not [21, 38], or only observe loss of MYH1 and MYH4[34]. These studies are typically limited to a single time points. Despite increased expression of less mature type IIb fiber genes, our data do not support a fiber type shift. We observe no progressive increase in oxidative or decrease in glycolytic genes, nor a shift in oxidative gene expression across myonuclear subpopulations. Instead, expression patterns are binary (high vs. low), not gradient. Based on protein expression, MYH4 appears stable while MYH2 and MYH1 are increased. This prompted us to consider new fiber formation, yet we see no rise in proliferative, embryonic, or myogenesis markers, consistent with prior findings highlighting no changes in embryonic protein expression[39]. Others have reported increased muscle damage[40], central nuclei[39], fibrosis[41], and impaired regeneration[40, 42] in cancer cachexia. However, we do not observe this, further suggesting that the increase in Myh1 and Myh2 is not derived from new or regenerating fibers. Although muscle fibers can co-express multiple MYH isoforms (hybrid fibers)[43], our immunofluorescence data do not indicate a notable increase in hybrid fibers during cachexia. Given this, we conclude that the increase in Myh1 and Myh2 expression reflects myofiber dedifferentiation which has been reported following muscle injury[44] and limb regeneration[45].

Extensive research has demonstrated that cancer cachexia disrupts skeletal muscle protein homeostasis by promoting a chronic catabolic state. Although these catabolic pathways are transcriptionally upregulated during cachexia progression[15, 16], it is unclear whether such transcriptional changes are confined to myonuclei. Recent snRNAseq studies in cachectic mice have identified distinct sub-populations of myonuclei expressing catabolic genes[19, 20]. Our results are consistent with these findings because we identified a PDAC-associated myonuclei sub-population that highly expresses cachexia-related genes, even prior to muscle wasting. However, we also observed that cachexia gene expression was not restricted solely to this PDAC-specific myonuclear subpopulation and did not overlap with *Myh1* and *Myh2* expressing myonuclei. This suggests that the cachexia gene signature is robustly upregulated prior to overt wasting, is ubiquitously expressed, and the shift in myosin expression modestly overlaps with cachexia genes highlighting that PDAC cachexia elicits myonuclear distinct transcriptional responses. Importantly, analysis of skeletal muscle from PDAC patients revealed a reduction in mature myosin isoform expression that did not transcriptionally overlap with regions of elevated cachexia gene expression, further supporting the existence of discrete myonuclear responses during cachexia progression.

In summary, pancreatic cancer induces population-specific switching of myosin isoforms and discrete activation of cachexia-related genes in skeletal muscle in both mice and human patients. Our findings implicate the loss of the tumor-induced muscle differentiation factor *Maf* and its downstream target *Myh4*, with a parallel gain in *Myh1* and *Myh2* expression during the progression of cachexia. These results reveal population-specific heterogeneity in cachexia gene activation, rather than a uniform upregulation of cachexia mediators across muscle tissue, likely contributing to the challenge of effectively targeting skeletal muscle wasting in cancer cachexia. We hypothesize that the fate of myonuclei occurs early in cachexia. We purpose that myonuclei either (1) don’t respond to systemic inflammation, (2) myonuclei express a cachexia or a dedifferentiation gene signature, or (3) myonuclei exhibit both a cachectic and dedifferentiation gene signature prompting irreversible changes prior to overt muscle wasting (graphical abstract). Future studies should focus on mapping the temporal dynamics and regulatory mechanisms driving these population-specific transcriptional changes to identify precise molecular targets for therapeutic intervention.

## Acknowledgments

This work was supported by P01 CA236778-01A1 (Zimmers, Koniaris, Ostrowski, Guttridge, Cao), R01CA257452 (Zimmers), I01 CX002046 (Zimmers), T32 CA254888 (Postdoctoral fellow Counts), NIH/NCI 1L70CA284463-01 (Counts). We thank the Histopathology Shared Resource for pathology studies, which is supported in part by the University Shared Resource Program at Oregon Health & Science University and the Knight Cancer Institute (P30 CA069533). We acknowledge expert technical assistance by staff in the Advanced Multiscale Microscopy Shared Resource, supported by the OHSU Knight Cancer Institute (NIH P30 CA069533). Equipment purchases included support by the OHSU Center for Spatial Systems Biomedicine, the MJ Murdock Charitable Trust, and the Collins Foundation. Sequencing was carried out by the Center for Medical Genomics using the Indiana University School of Medicine Flow Cytometry Core, and the Immunohistochemistry Research Core (NIH grant P30 CA082709).

---

**Supplemental Figure 1: Single nucleus filtering and cell type identification.**

**A)** Gastrocnemius (GAS) and tibialis anterior (TA) mass relative to initial body weight. **B)** Table of absolute muscle mass at the time of tissue harvest. One-way ANOVA using Tukey’s multiple comparison tests. *Different from sham, $different from D6, #different from D9, ^different from D12. P<0.05 all. **C)** Single nucleus filtering of features (200-5000 genes) and **D)** Mitochondria counts (<5%). **E)** Unbiased t-SNE plot of all clusters in the integrated sham and PDAC conditions. **F)** Cell type gene expression and percentage of nuclei expressing the genes in the unbiased clustering.

**Supplemental Figure 2: Myonuclei sub-cluster identification and bulk RNA sequencing expression of skeletal muscle myosin and *Maf* genes in PDAC cachexia.**

**A)** Heat map of Z-scores for the top 5 biomarkers of each myonuclei sub-population. **B)** Skeletal muscle developmental and maturation gene expression and percentage of nuclei expressing the genes. **C)** Myosin heavy chain gene isoforms expressed as normalized counts. **D)** Myosin light chain gene isoforms expressed as normalized counts. **E)** Transcription factors enriched in fully differentiated fibers from Dos Santos et al. [8] during body weight loss in PDAC cachexia. **F)** Large and small *Maf* genes expressed as normalized counts. (**C-E)** One-way ANOVA with Tukey’s post hoc or Kruskal-Wallis (only used in D Myl4) test with Dunn’s multiple comparison post hoc test was performed. *p<0.05. **F)** Heat map of known *Maf* targets in the snRNAseq data displayed at the pseudo-bulk (all cell types) and myonuclei level, and bulk RNAseq gene expression across the time course comparing PDAC vs sham. Data are expressed as fold change. **G)** Associations of body weight change from pre, accounting for tumor mass, with myosin heavy chain and light chain isoforms, and *Maf* targets. Statistical test: Pearson correlation coefficient. **H)** Heat map of top 100 cachexia genes with differentiated muscle genes. Statistical test: Pearson correlation coefficient from bulk RNAseq data in both Sham and PDAC mice. Associations in Table 7. **I)** Association of sum differentiation gene score with sum cachexia gene score. Gene ranks found in Figure 4H.

**Supplemental Figure 3: Fast myosin heavy chain protein expression and *Maf* gene expression during PDAC cancer cachexia.**

**A)** Immunofluorescence of MYH4 and MYH2 and RNAscope of *Maf* in the sham and PDAC conditions. **B)** Immunofluorescence of MYH1 in the sham and PDAC conditions. Immunofluorescence was completed in longitudinal sections of the gastrocnemius. We first performed MYH7 expression to identify fibers not expressing MYH7 to prevent bias by imaging the more oxidative portion of the gastrocnemius muscle.

**Supplemental Figure 4: PDAC alters oxidative phenotypes in a sub-group specific fashion, and lack of evidence to support a fiber type shift in the myonuclei by snRNAseq nor in muscle by bulk RNAseq.**

**A)** Top 3 GO biological processes in myonuclei sub-populations. Enrichment score graphed on the x-axis. FDR is expressed by bubble color, and total genes in the biological pathway expressed by bubble size. Publicly available Enrichr was used to determine the top biological pathways from upregulated genes. **B)** Trajectory analysis of myonuclei sub-populations in the sham and PDAC conditions. Initial starting point (1) was performed using unbiased identification of the integrated samples. **C-E)** Gene expression and percentage of nuclei expressing those genes from snRNAseq. Genes are grouped by those increased in oxidative fibers **(C),** increased glycolytic fibers **(D)**, or identified fiber type switching **(E). F-H)** Gene expression of oxidative **(F)**, glycolytic **(G),** and fiber type switching **(H)** genes across the time of PDAC in bulk RNAseq. Data is expressed as fold change of PDAC vs sham across the time course. Sham transcript counts are listed in the last column.

**Supplemental Figure 5: Low evidence for PDAC effects on regeneration, fibrosis, or proliferation.**

**A-C)** Heat maps of skeletal muscle embryonic **(A),** proliferative **(B)**, and fibrillogenesis and myogenesis **(C)** genes in bulk RNAseq across the time course comparing PDAC vs sham. Data are expressed as fold change. **D)** H&E stains of tibialis anterior (TA) cross-sections in the sham and PDAC conditions. Yellow arrows identify centrally located nuclei. **E)** Trichrome stains of TA cross-sections in the sham and PDAC conditions.

**Supplemental Figure 6: Single Nucleus cell counts for mouse and human skeletal muscle in pancreatic cancer cachexia.**

**A-C)** Murine nuclei counts pre and post filtering, in each cell type population and in each myonuclei sub-population. **D-E)** Human patient nuclei count pre and post filtering, and each cell type population.

## Data Availability

All sequencing data files and codes are publicly available. The raw murine muscle single nucleus RNAseq and muscle bulk RNAseq data generated in this study are publicly available through NCBI-GEO (GSE297184). Partek and Python coding files can be found in Supplemental Figure 8.

## Notes

### Competing Interest Statement

TAZ has been compensated for consulting work on cancer cachexia, has carried out sponsored research for Leap Therapeutics, and was/is a member of the Scientific Advisory Board of Emmyon, Inc. and PeleOs. However, none of these financial relationships concern the research presented here. The authors declare no further conflicts of interest.

