## Supplementary Data for "Pancreatic Cancer Induces Population-Specific Switching of Myosin Isoforms and Discrete Activation of Cachexia Genes in Skeletal Muscle Myocytes"

Graphical Abstract

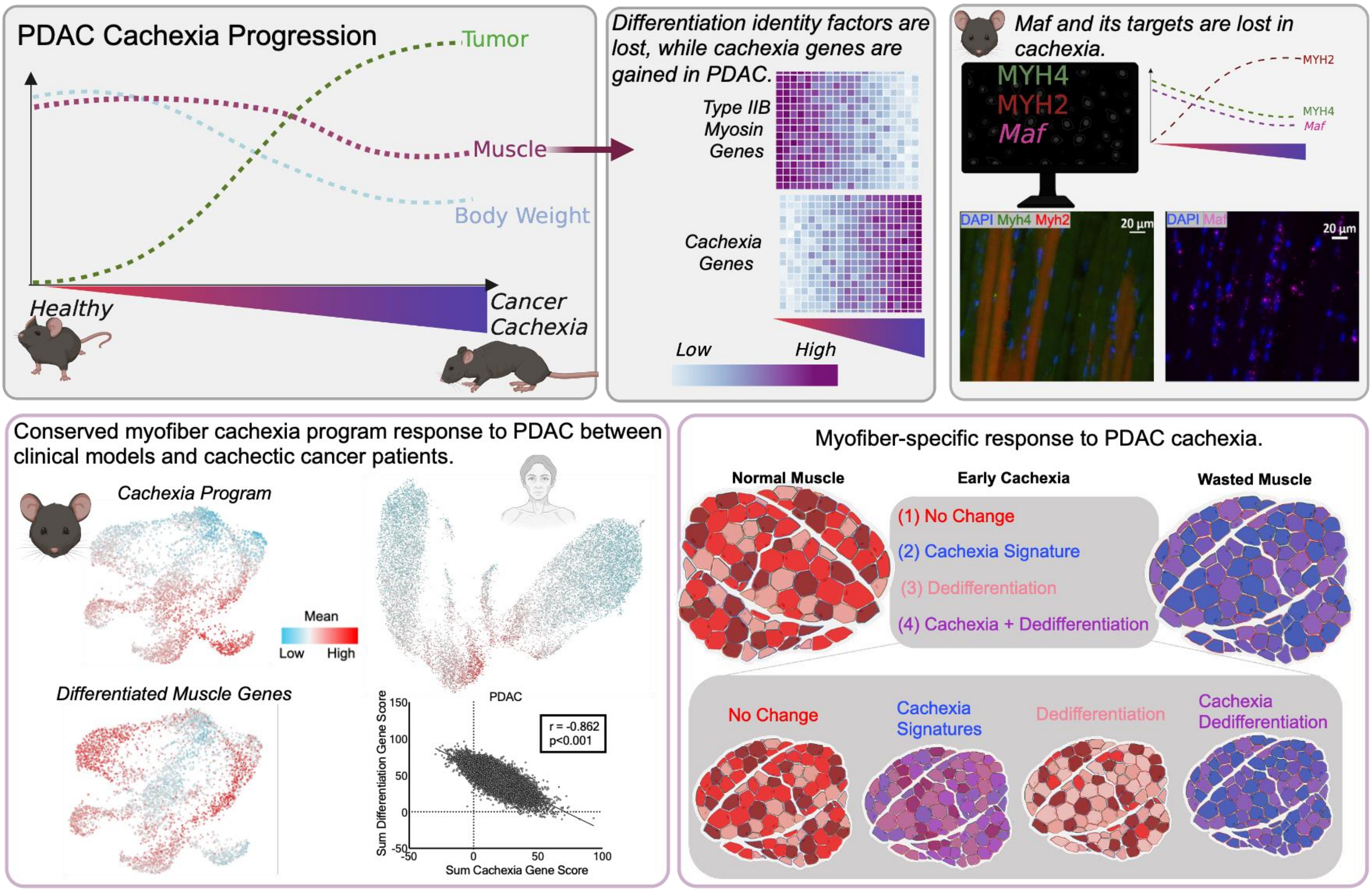


We observe a progressive loss of muscle differentiation factor *Maf* and its target *Myh4*, accompanied by increased expression of *Myh1* and *Myh2* during cachexia progression. We uncover population-specific heterogeneity in the activation of genes associated with cachexia, rather than a uniform upregulation of cachexia mediators across muscle tissue. Impactfully, the loss of muscle identity and the gain of catabolic gene expression occurred in distinct myonuclear populations. Translating these findings to humans, we observe a similar myonuclear response in patients with PDAC cachexia, underscoring a conserved myofiber response between clinical models and cachectic cancer patients. Our data suggest that the fate of myonuclei occurs prior to overt muscle wasting, when cachexia gene expression only modestly overlaps with differentiation factors, with a strong association after irreversible muscle wasting.

Supplemental Figure 1: Single nucleus filtering and cell type identification.

A.

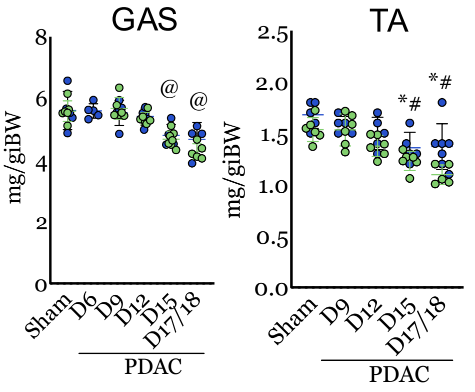


B.

| | Sham | PDAC D6 | PDAC D9 | PDAC D12 | PDAC D15 | PDAC D17/18 |
|---|---|---|---|---|---|---|
| N | 11 | 5 | 11 | 11 | 12 | 9 |
| GAS (mg) | 148 (18) | 156(3) | 146(12) | 140(8) | 128(14)*^ | 118(15)*^# |
| TA (mg) | 42(6) | 43(4) | 40(4) | 38(4) | 35(4)* | 33(6)* |
| QUAD (mg) | 199(32) | 194(15) | 182(30) | 184(17) | 175(26) | 154(24)* |

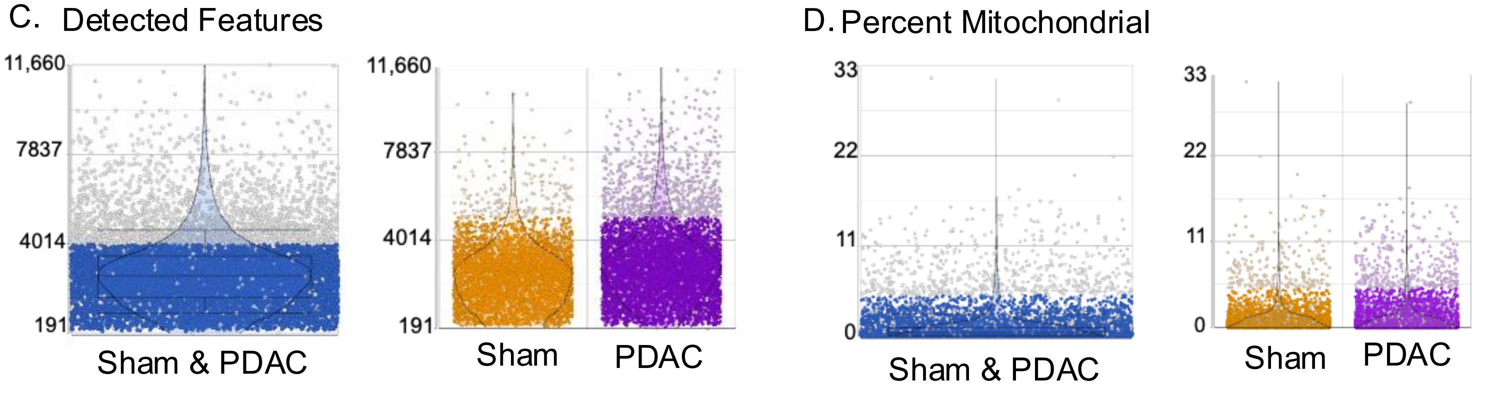


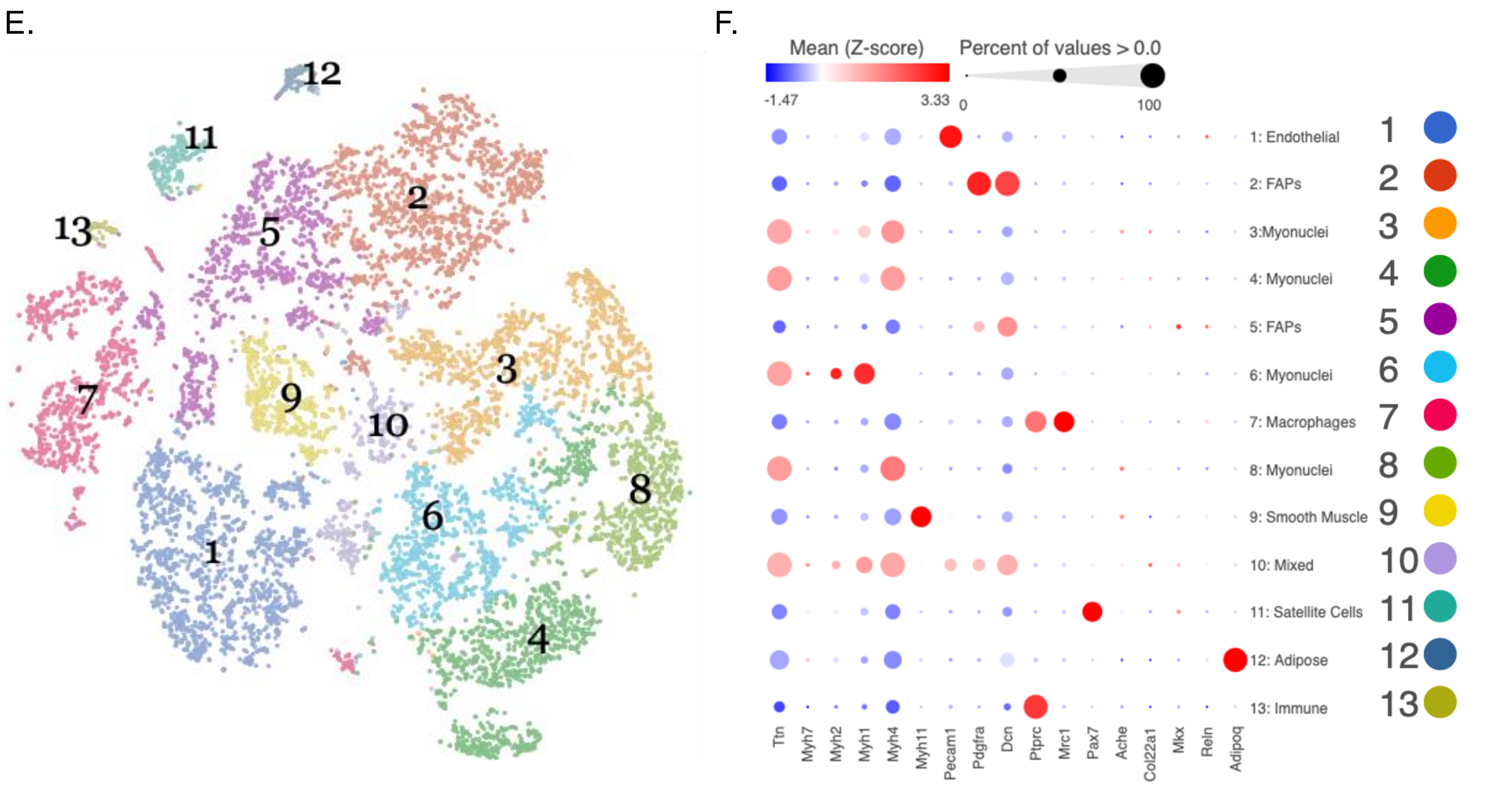

Supplemental Figure 2: Myonuclei sub-cluster identification and bulk RNA sequencing expression of skeletal muscle myosin and Maf gene isoforms in PDAC cachexia.

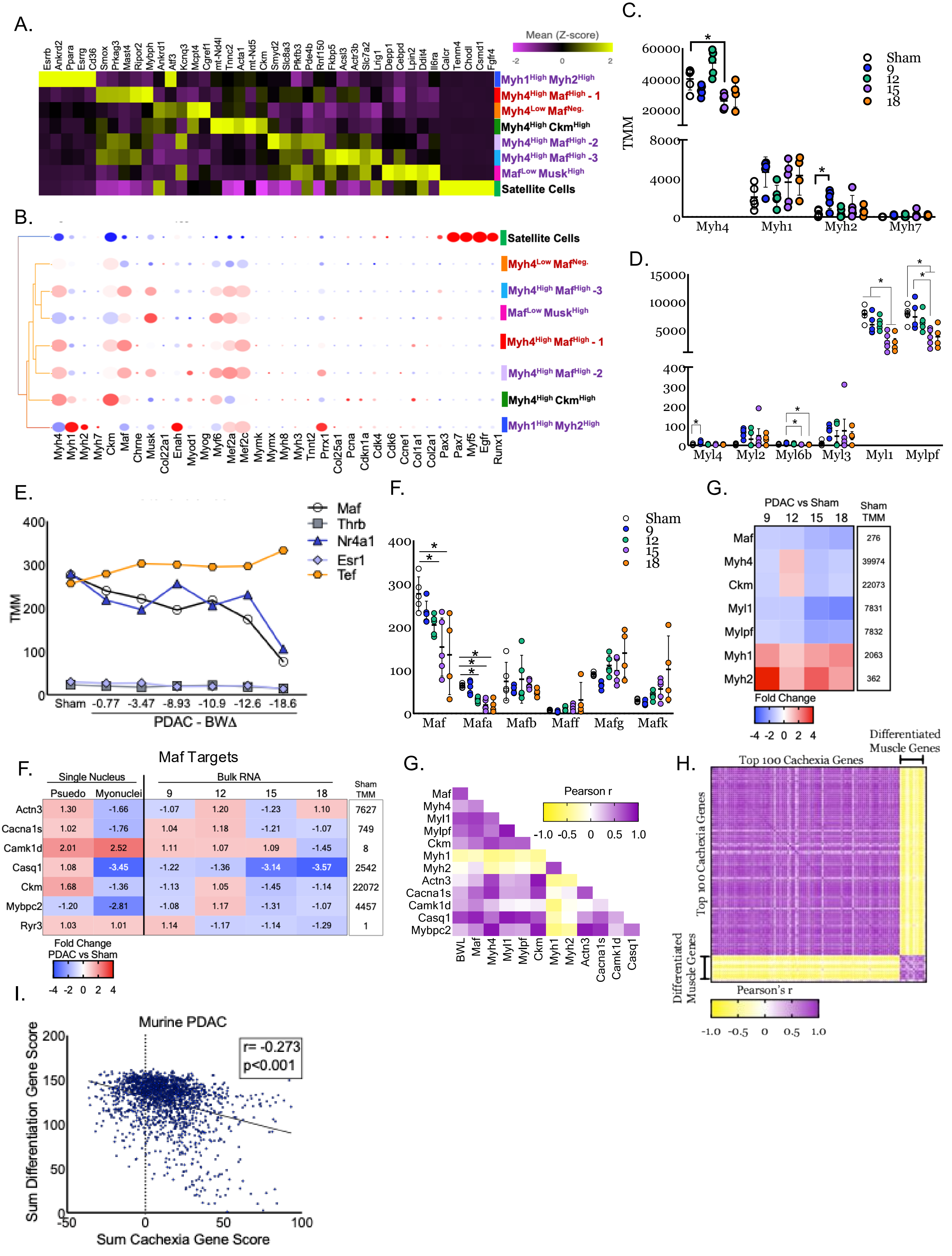


| | PDAC vs Sham: 9 | 12 | 15 | 18 | Sham TMM |
|---|---|---|---|---|---|
| Maf | | | | | 276 |
| Myh4 | | | | | 39974 |
| Ckm | | | | | 22073 |
| Myl1 | | | | | 7831 |
| Mylpf | | | | | 7832 |
| Myh1 | | | | | 2063 |
| Myh2 | | | | | 362 |

| | Single Nucleus: Psuedo | Myonuclei | Bulk RNA: 9 | 12 | 15 | 18 | Sham TMM |
|---|---|---|---|---|---|---|---|
| Actn3 | 1.30 | -1.66 | -1.07 | 1.20 | -1.23 | 1.10 | 7627 |
| Cacna1s | 1.02 | -1.76 | 1.04 | 1.18 | -1.21 | -1.07 | 749 |
| Camk1d | 2.01 | 2.52 | 1.11 | 1.07 | 1.09 | -1.45 | 8 |
| Casq1 | 1.08 | -3.45 | -1.22 | -1.36 | -3.14 | -3.57 | 2542 |
| Ckm | 1.68 | -1.36 | -1.13 | 1.05 | -1.45 | -1.14 | 22072 |
| Mybpc2 | -1.20 | -2.81 | -1.08 | 1.17 | -1.31 | -1.07 | 4457 |
| Ryr3 | 1.03 | 1.01 | 1.14 | -1.17 | -1.14 | -1.29 | 1 |

Supplemental Figure 3: Fast myosin heavy chain protein expression and *Maf* gene expression during PDAC cancer cachexia.

A.

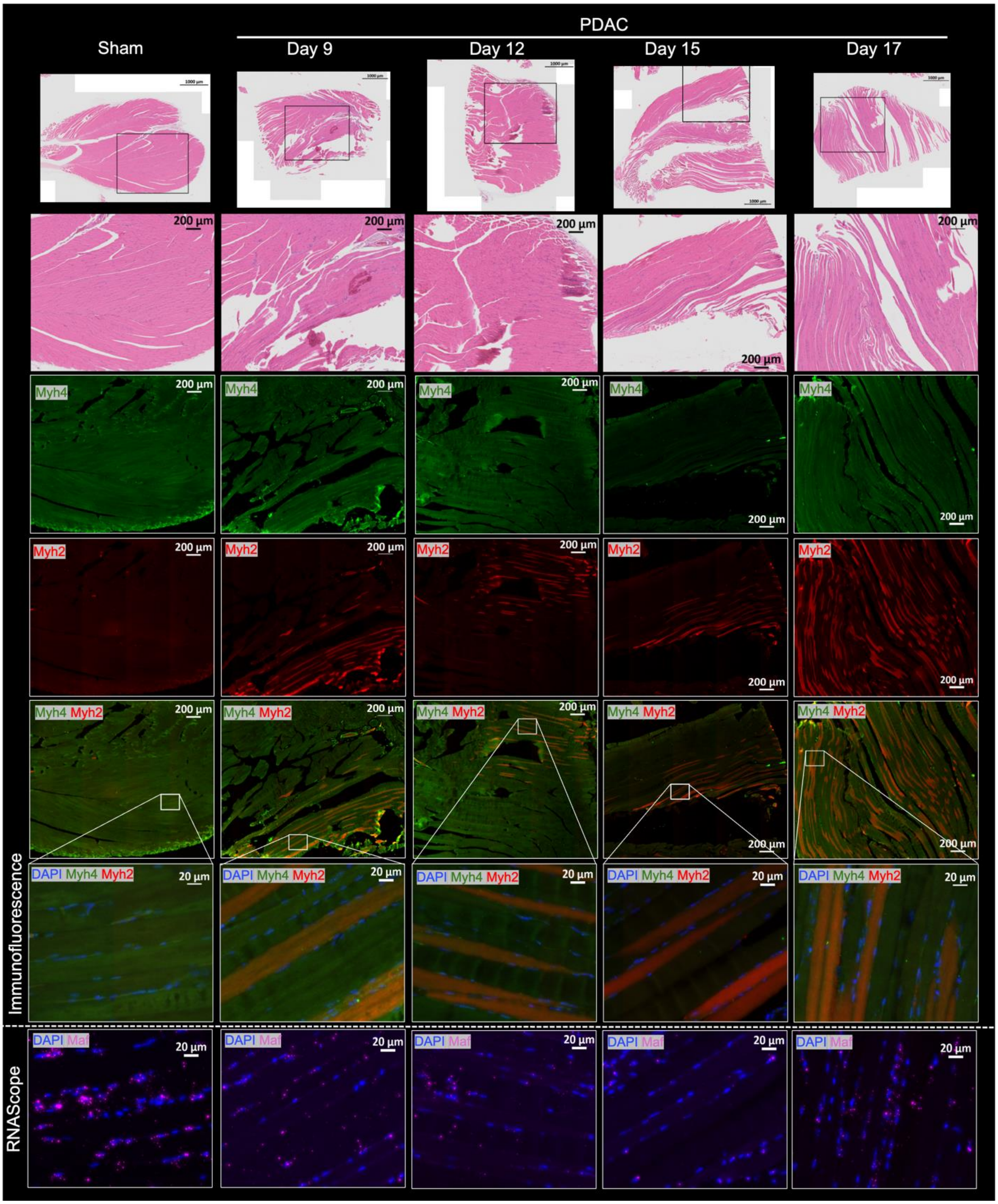


B.

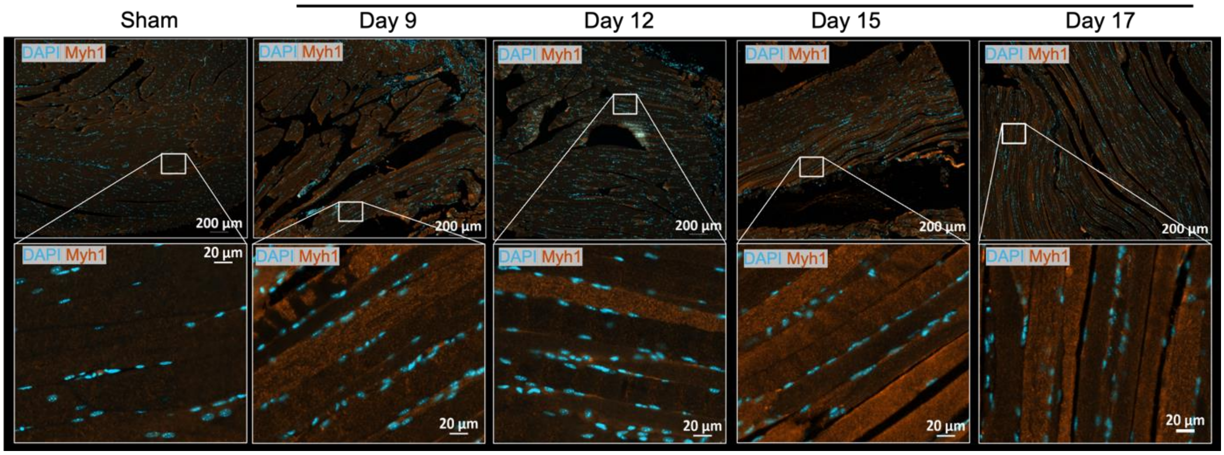

Supplemental Figure 4: PDAC alters oxidative phenotypes in a subgroup-specific fashion, and lack of evidence to support a fiber type shift in the myonuclei by snRNAseq nor in muscle by bulk RNAseq.

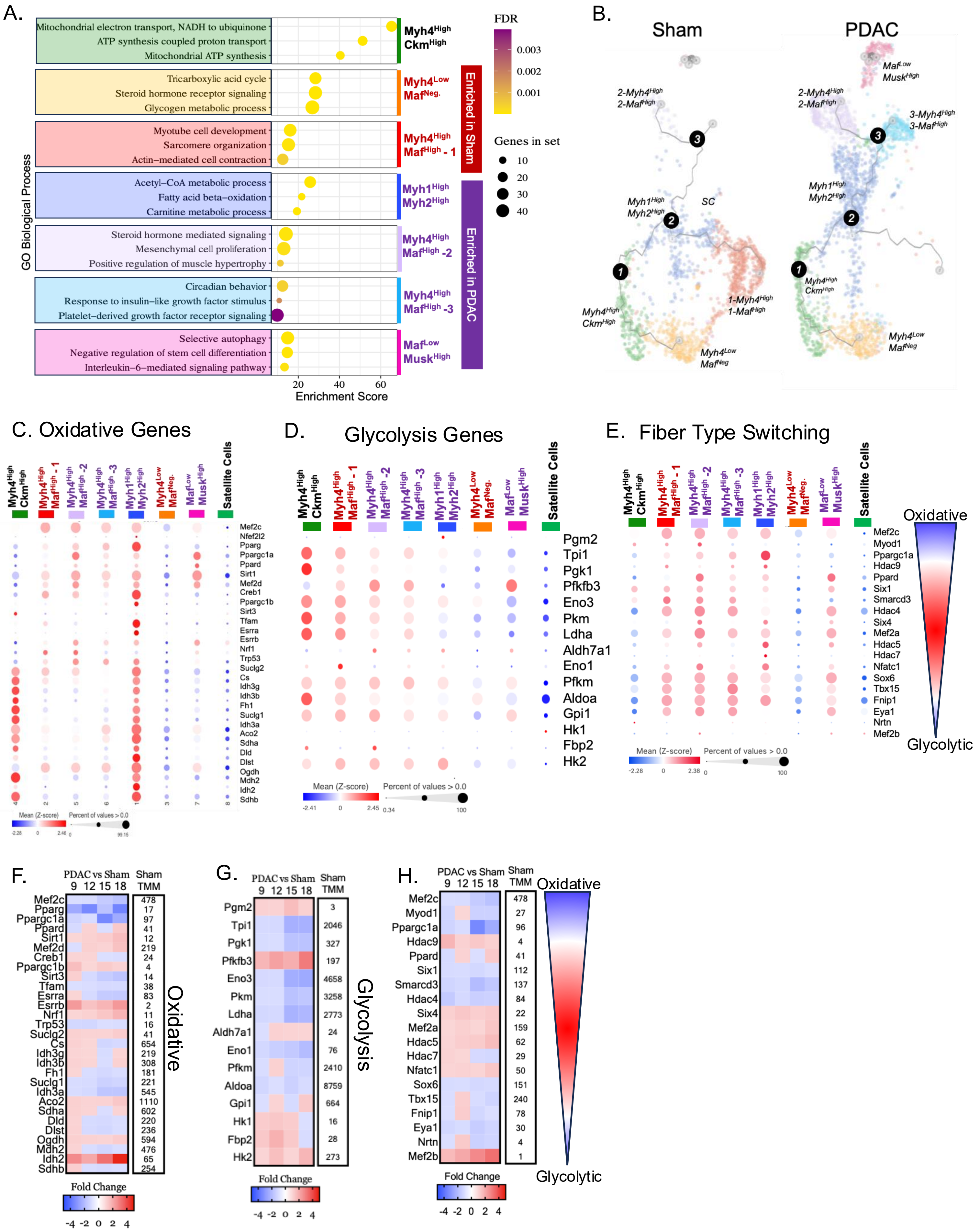

Supplemental Figure 5: Low evidence for PDAC effects on regeneration, fibrosis, or proliferation.

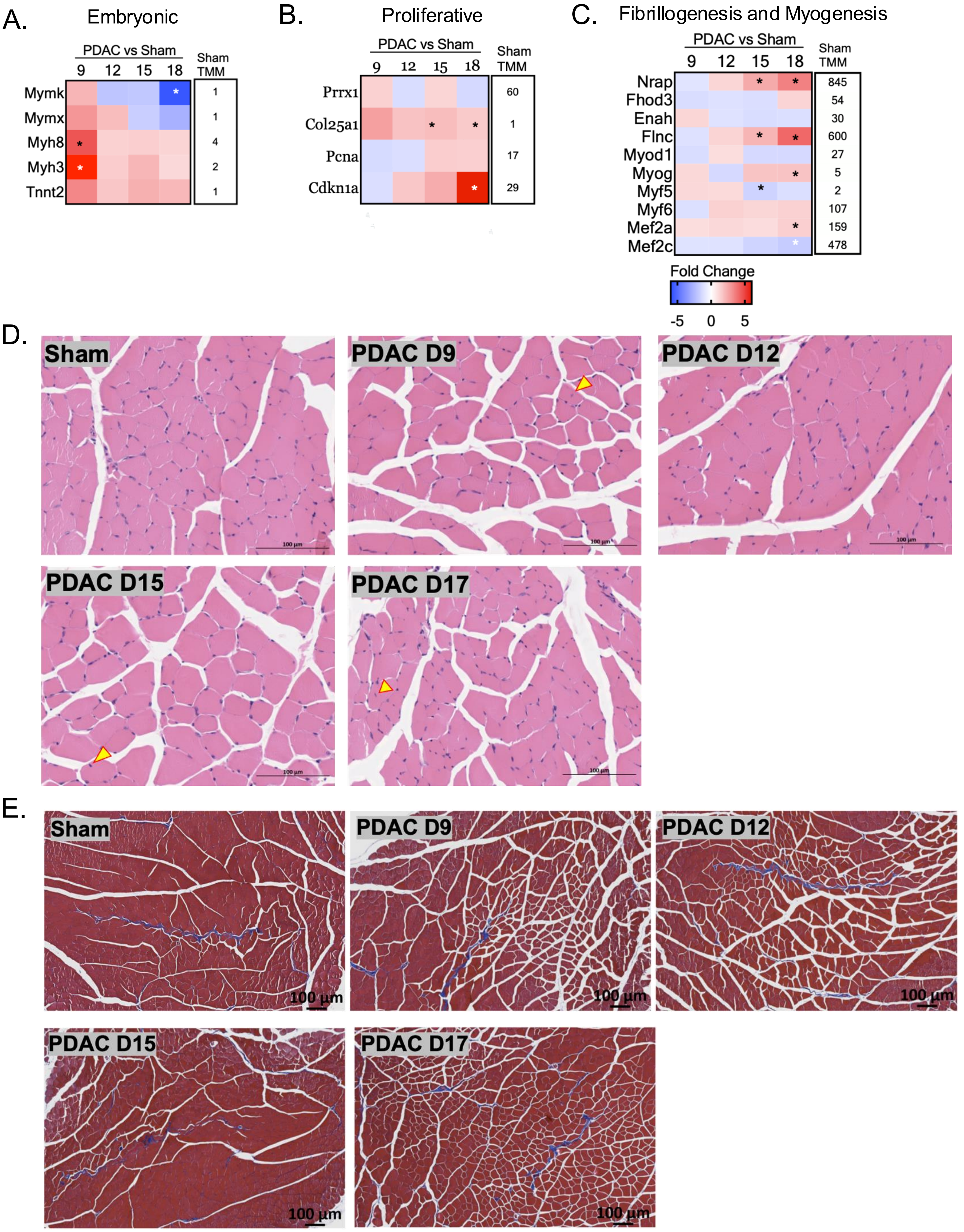

Supplemental Figure 6: Single Nucleus cell counts for mouse and human skeletal muscle in pancreatic cancer cachexia.

A.

| | Sham & PDAC | Sham | PDAC |
|---|---|---|---|
| Pre-Filtering Nuclei (#) | 12,573 | 4898 | 7675 |
| Post-Filtering Nuclei (#) | 11,083 | 4497 | 6586 |
| % Used for Analysis | 87.1% | 91.8% | 85.8% |

B.

| Cell Type | Sham & PDAC | Sham | PDAC |
|---|---|---|---|
| Adipocytes (#) | 104 | 29 | 75 |
| Endothelial Cells (#) | 1583 | 633 | 950 |
| FAPs (#) | 2810 | 1271 | 1539 |
| Immune (#) | 67 | 42 | 25 |
| Macrophages (#) | 995 | 446 | 549 |
| Mixed Cells (#) | 459 | 111 | 348 |
| Myonuclei (#) | 4335 | 1684 | 2651 |
| Satellite Cells (#) | 210 | 82 | 128 |
| Smooth Muscle (#) | 520 | 199 | 321 |

C.

| Cluster # | Cell Type | Sham & PDAC | Sham | PDAC |
|---|---|---|---|---|
| 1 | Myh1$^{High}$, Myh2$^{High}$ (#) | 1107 | 324 | 783 |
| 2 | Myh4$^{High}$, Maf$^{High}$ – 1 (#) | 708 | 699 | 9 |
| 3 | Myh4$^{Low}$, Maf$^{Neg.}$ (#) | 686 | 344 | 342 |
| 4 | Myh4$^{High}$, Ckm$^{High}$ (#) | 642 | 260 | 382 |
| 5 | Myh4$^{High}$, Maf$^{High}$ -2 (#) | 553 | 53 | 500 |
| 6 | Myh4$^{High}$, Maf$^{High}$ -3 (#) | 349 | 1 | 348 |
| 7 | Maf$^{Low}$, Musk$^{High}$ (#) | 291 | 4 | 287 |
| 8 | Satellite Cells (#) | 209 | 81 | 128 |

D.

| | Pancreatic Cancer Patient |
|---|---|
| Pre-Filtering Nuclei # | 15,307 |
| Post-Filtering Nuclei # | 15,079 |
| % Remained after filtering | 98.5% |

E.

| Cell Type (Nuclei #) | Pancreatic Cancer Patient |
|---|---|
| Myonuclei (#) | 14203 |
| FAPs (#) | 645 |
| Endothelial & Adipocytes (#) | 153 |
| Immune Cells (#) | 40 |
| MuSC (#) | 38 |

Extended Methods:

**Mouse Muscle Single Nucleus Isolation and Analysis:**

*Nuclei Isolation:* We isolated nuclei from the quadriceps muscle of 4 sham mice and 4 PDAC D12 mice. Approximately 50mg of flash-frozen quadriceps muscle tissue was powdered in liquid nitrogen using a mortar and pestle. Samples were then dounced on ice in 7ml of NPB-VRC (0.25M sucrose, 0.01M Hepes pH7.5, 0.01M KCL, 0.1%NP40, 0.01M VRC added at RT, 1X halt protease inhibitor, 0.001M DTT and 1X RNase inhibitor) 10 times. Samples sat on ice for 5mins, then strained through a 100um strainer into a 50ml tube. Samples were spun at 200xg at 4°C for 10mins in a swinging bucket centrifuge. The supernatant was removed and 1ml of PBS-N-VRC (PBS-N: 0.10% NP40 and 1X PBS Mg and Ca free, VRC: 0.005M VRC at RT) was added to gently disturb the pellet. Next, nuclei were resuspended in 4ml of PBS-N-VRC. Samples underwent a second spin of 200xg at 4°C for 5mins and were resuspended in 1000µl of PBS-N-VRC, then strained through a 40µm filter. Nuclei were FACS sorted using DAPI, and we collected ~500,000 nuclei/sample. After sorting, samples were suspended in 1000µl of PBS-N-VRC and spun 200xg for 10 minutes. Bubbles were removed from the tubes and lid; supernatant was slowly poured out, and samples were resuspended in 30µl of PBS-N. Samples were counted at the IU School of Medicine Genomics Core; the sham and PDAC D12 samples were resuspended, pooled, and submitted for sequencing.

*Sequencing:* Single-nuclei 3' RNA-seq assay was conducted using the 10x Chromium single-cell system (10x Genomics, Inc). Each single-nuclei suspension was first checked for nuclei quality and quantity. The four samples were pooled for each condition. Around 16,500 nuclei from each condition were loaded into one well of a multiple-channel microfluidics chip with 10,000 targeted cells. Single-cell gel beads in an emulsion containing barcoded oligonucleotides and reverse transcriptase reagents were generated with the Next Gem single-cell reagent kit (10X Genomics). cDNA was synthesized and amplified. The quality of cDNA and library were examined by Bioanalyzer at each step. The final dual-indexed library was sequenced on an Illumina NovaSeq 6000. 100-bp reads, including cell barcode and UMI sequences were generated.

*Filtering and analysis:* 10X chromium fastq files were uploaded to Partek Flow. First, we filtered by excluding nuclei with less than 200 or more than 5000 genes, and nuclei with more than 5% of mitochondria (Supplemental Figure 1C,D). Then, counts were normalized to counts per million, and genes with a value of 0 in 100% of the nuclei were removed. See supplemental text for Partek command lines (Supplemental Figure 8). We generated an unbiased PCA to identify the number of PCs to include for downstream analysis that will encompass >99% of the variance. Cell types were identified based on the expression of validated cell-type genes (Supplemental Figure 1E). Similar cell types were grouped, and then cell type biomarkers were generated (Figure 1G, Supplemental Table 1). Hurdle analysis was used to compare cancer vs non-cancer, and myonuclei sub-populations. Gene enrichment score from the biological process was determined using genes with FDR <0.05 from cluster comparisons, fold change greater than 1, and the total genes in the set are less than 50.

**Mouse Muscle Bulk RNA sequencing cDNA library preparation, sequencing, and filtering***:* cDNA library preparation includes mRNA purification/enrichment, RNA fragmentation, cDNA synthesis, and ligation of index adaptors using the KAPA mRNA Hyperprep Kit prep (cat: KK8581). Each resulting indexed library was quantified, and its quality was assessed by NovaSeq 6000. 5μL of 2nM pooled libraries per lane were denatured, neutralized, and applied to the cBot for flow cell deposition and cluster amplification before loading to 100bp paired-end sequencing (Illumina, Inc). Approximately 30-50 million reads per library were generated. Only reads with Phred quality score of 35 or more were used for used to measure sequencing quality. Reads were mapped to the mm10 reference genome using STAR. We performed noise reduction filtering and performed DESeq2. We performed one-way ANOVA on the trimmed mean of M values (TMM) normalized counts.

**Human Muscle Single Nucleus Isolation:**

Method of isolation: Sample reagents and materials were pre-chilled and kept on ice as much as possible during the nuclei isolation steps. All centrifugation steps were carried out at 4°C using a swing bucket rotor for 5 minutes: 500g, acceleration 9, deceleration 7. 50 mg flash frozen muscle stored at -80 was used for source material. 750 µl TsT nuclei lysis buffer ((1X salt-Tris (sT) buffer (146 mM NaCl, 10 mM Tris-HCl pH 7.5, 1 mM $CaCl_2$, 21 mM $MgCl_2$, 250 mM Sucrose, 1X Halt, 800U/ml RNase Inhibitor, 1mM DTT) + 0.03% Tween-20)) was added to 50 mg of flash frozen tissue in a 1.7 ml tube. Scissors were used to finely chop the tissue for about 5-10 minutes. The minced tissue was transferred to a pre-chilled dounce homogenization tube. 750 µl additional TsT buffer was used to collect all minced tissue. Pre-chilled pestles (placed in dounce buffer on ice) were used to dounce homogenize the tissue. Dounce homogenization was done using 20 strokes each using pestle A followed by pestle B. Homogenate was transferred to a pre-chilled 5 ml round bottom tube, and an additional 500 µl TsT was used to wash out the dounce and collect all of the homogenate. A wide-bore pipet tip was used to pipette the homogenate up and down a few times to mix. Homogenates were chilled on ice for 5 minutes then filtered through a 40 µm cell strainer into a 50 ml conical tube. Homogenate was washed through with an additional 500 µl TsT buffer. Filtered sample was transferred into a pre-chilled 15 ml conical tube and centrifuged for 5 minutes. Supernatant was removed leaving about 50 µl in the bottom of the tube with the nuclei pellet. Pellet was resuspended in 1 ml 1X sT buffer (no Tween-20. Sample was incubated on ice then centrifuged 5 minutes. Supernatant was removed, and the pellet was resuspended in 1 ml 1X Fluent NWB containing RNase inhibitor. A 1ml narrow bore pipet tip was used to mix the sample prior to the addition of 4ml additional NWB was added with a serological pipette on ice. A 10 µm uberstrainer with a 5 ml luer-lok syringe attached was assembled to a 50 ml conical tube on ice. 2 ml of nuclei suspension was added to the uberstrainer, and negative pressure was applied using the 5 ml syringe (less than 1ml/5s) and not exceeding the 3 ml mark. If necessary, the syringe was removed, reset to 0, and repeated. After filtering the initial 2 ml of suspension, the remaining sample was added and filtered. The sample was transferred to a 15 ml conical tube and centrifuged for 5 minutes. Supernatant was removed and nuclei were resuspened in 200 µl NSB (1X NSB, 0.8% BSA, 0.5U/µl RNase Inhibitor). Nuclei quality was assessed visually and quantitatively using trypan blue, Hoescht, and the Invitrogen EVOs scope and Countess II FL. Fresh nuclei were resuspended (3400/ µl) in NSB and used immediately for library preparation using the Fluent PIPseq V T10 3' Single Cell RNA Kit according to the manufacturers protocol. Library QC and sequencing was done by Novogene using a NovaSeq X Plus Series.

**Supplemental Figure 8: snRNAseq Partek Command Lines**

**Murine**
**Filter Cells**
java: setup
/opt/partek_flow/bin/tabular/flow_table --threads 0 -p /home/flow/FlowData/userdata/bcounts/Project_snRNA Seq Trial_Skeletal Muscl_7/FilterCellsResult-24942551988630052/parameters1606283719335478044l.json
java: setup
java: setup
/opt/partek_flow/bin/datablock/filter_datablock --input /home/flow/FlowData/userdata/bcounts/Project_snRNA Seq Trial_Skeletal Muscl_7/FilterCellsResult-24942551988630052/input2729629437687146888.json --output /home/flow/FlowData/userdata/bcounts/Project_snRNA Seq Trial_Skeletal Muscl_7/FilterCellsResult-24942551988630052/output2945392671468409883.json --row-vector-filter /home/flow/FlowData/userdata/bcounts/Project_snRNA Seq Trial_Skeletal Muscl_7/FilterCellsResult-24942551988630052/rowIndices1833413288996l228933.bin --sort_indices --threads 0
java: setup
java: java
java: setup
java: setup
/opt/partek_flow/bin/datablock/datablock_stats --input /home/flow/FlowData/userdata/bcounts/Project_snRNA Seq Trial_Skeletal Muscl_7/FilterCellsResult-24942551988630052/filtered.mat.matrix --sample-ids /home/flow/FlowData/userdata/bcounts/Project_snRNA Seq Trial_Skeletal Muscl_7/FilterCellsResult-24942551988630052/filtered.observations --output /home/flow/FlowData/userdata/bcounts/Project_snRNA Seq Trial_Skeletal Muscl_7/FilterCellsResult-24942551988630052/feature_distribution.txt --threads 0
java: java
java: java

**Normalization**
/opt/partek_flow/bin/datablock/normalize_datablock --threads 0 --input /home/flow/FlowData/userdata/bcounts/Project_snRNA Seq Trial_Skeletal Muscl_7/NormalizationResult-24942743465943509/input8183151120937869412.json --output /home/flow/FlowData/userdata/bcounts/Project_snRNA Seq Trial_Skeletal Muscl_7/NormalizationResult-24942743465943509/normalized.mat.matrix --transform-on Cells --total-count --add 1.0 --log 2.0 --output_effective_libsize /home/flow/FlowData/userdata/bcounts/Project_snRNA Seq Trial_Skeletal Muscl_7/NormalizationResult-24942743465943509/normalized.effective_lib_sizes.txt
java: setup
java: java
java: setup
java: setup
java: if
/opt/partek_flow/bin/quantification/generate_expression_chart_data --input /home/flow/FlowData/userdata/bcounts/Project_snRNA Seq Trial_Skeletal Muscl_7/NormalizationResult-24942743465943509/normalized.mat.matrix --sample-ids /home/flow/FlowData/userdata/bcounts/Project_snRNA Seq Trial_Skeletal Muscl_7/FilterCellsResult-24942551988630052/filtered.observations --chart-output /home/flow/FlowData/userdata/bcounts/Project_snRNA Seq Trial_Skeletal Muscl_7/NormalizationResult-24942743465943509/expressionChartCache.json --stats-output /home/flow/FlowData/userdata/bcounts/Project_snRNA Seq Trial_Skeletal Muscl_7/NormalizationResult-24942743465943509/feature_distribution.txt --threads 4
/opt/partek_flow/bin/quantification/generate_expression_chart_data --input /home/flow/FlowData/userdata/bcounts/Project_snRNA Seq Trial_Skeletal Muscl_7/FilterCellsResult-24942551988630052/filtered.mat.matrix --sample-ids /home/flow/FlowData/userdata/bcounts/Project_snRNA Seq Trial_Skeletal Muscl_7/FilterCellsResult-24942551988630052/filtered.observations --chart-output /home/flow/FlowData/userdata/bcounts/Project_snRNA Seq Trial_Skeletal Muscl_7/FilterCellsResult-24942551988630052/expressionChartCache.json --threads 4
/opt/partek_flow/bin/datablock/datablock_stats --input /home/flow/FlowData/userdata/bcounts/Project_snRNA Seq Trial_Skeletal Muscl_7/FilterCellsResult-24942551988630052/filtered.mat.matrix --sample-ids /home/flow/FlowData/userdata/bcounts/Project_snRNA Seq Trial_Skeletal Muscl_7/FilterCellsResult-24942551988630052/filtered.observations --output /home/flow/FlowData/userdata/bcounts/Project_snRNA Seq Trial_Skeletal Muscl_7/NormalizationResult-24942743465943509/dataBlockStats3655861023786499198.txt --threads 0
java: java
java: setup
/opt/partek_flow/bin/datablock/datablock_stats --input /home/flow/FlowData/userdata/bcounts/Project_snRNA Seq Trial_Skeletal Muscl_7/NormalizationResult-24942743465943509/normalized.mat.matrix --sample-ids /home/flow/FlowData/userdata/bcounts/Project_snRNA Seq Trial_Skeletal Muscl_7/FilterCellsResult-24942551988630052/filtered.observations --calculate-percentiles --output /home/flow/FlowData/userdata/bcounts/Project_snRNA Seq Trial_Skeletal Muscl_7/NormalizationResult-24942743465943509/feature_distribution.txt --threads 4

java: java

**Filter**
/opt/partek_flow/bin/quantification/find_feature_indices --input /home/flow/FlowData/userdata/bcounts/Project_snRNA Seq Trial_Skeletal Muscl_7/NormalizationResult-24942743465943509/normalized.mat.matrix --output /home/flow/FlowData/userdata/bcounts/Project_snRNA Seq Trial_Skeletal Muscl_7/FeatureFilterResult-24943219446485782/filter13561189958042654435.txt --threads 0 --expression_method value --operation eq --expression_value 0.0 --expression_percent 100.0
java: setup
java: setup
/opt/partek_flow/bin/datablock/filter_datablock --input /home/flow/FlowData/userdata/bcounts/Project_snRNA Seq Trial_Skeletal Muscl_7/FeatureFilterResult-24943219446485782/input17937459919364486456.json --output /home/flow/FlowData/userdata/bcounts/Project_snRNA Seq Trial_Skeletal Muscl_7/FeatureFilterResult-24943219446485782/output8374997802192566233.json --column-filter /home/flow/FlowData/userdata/bcounts/Project_snRNA Seq Trial_Skeletal Muscl_7/FeatureFilterResult-24943219446485782/filter13561189958042654435.txt --sort_indices --threads 0
java: if
java: setup
java: setup
java: setup
/opt/partek_flow/bin/single_cell/sc_qa_qc --input-data /home/flow/FlowData/userdata/bcounts/Project_snRNA Seq Trial_Skeletal Muscl_7/FeatureFilterResult-24943219446485782/filtered.mat.matrix --input-features /home/flow/FlowData/userdata/bcounts/Project_snRNA Seq Trial_Skeletal Muscl_7/FeatureFilterResult-24943219446485782/filtered.col.matrix --input-samples /home/flow/FlowData/userdata/bcounts/Project_snRNA Seq Trial_Skeletal Muscl_7/FeatureFilterResult-24943219446485782/filtered.observations --input-sample-annotation /home/flow/FlowData/userdata/bcounts/Project_snRNA Seq Trial_Skeletal Muscl_7/FeatureFilterResult-24943219446485782/filtered.row.annotation --compute-iff-exists --output /home/flow/FlowData/userdata/bcounts/Project_snRNA Seq Trial_Skeletal Muscl_7/FeatureFilterResult-24943219446485782/scQAQC13681260807459158831.txt --threads 0 --input-annotation /home/flow/FlowData/library_files/user0/mm10/mm10_ensembl_release102_v2/Annotation file/mm10_ensembl_release102_v2.pannot
java: setup
/opt/partek_flow/bin/combine_datablocks/merge_attributes --threads 0 --specification /home/flow/FlowData/userdata/bcounts/Project_snRNA Seq Trial_Skeletal Muscl_7/FeatureFilterResult-24943219446485782/Merge_Specification2428296417012808514.txt --input /home/flow/FlowData/userdata/bcounts/Project_snRNA Seq Trial_Skeletal Muscl_7/FeatureFilterResult-24943219446485782/outputRowAnnotation1507554036790592818.txt --output /home/flow/FlowData/userdata/bcounts/Project_snRNA Seq Trial_Skeletal Muscl_7/FeatureFilterResult-24943219446485782/temp4242365981358455883.row.annotation
java: java
java: if
java: setup
/opt/partek_flow/bin/quantification/generate_expression_chart_data --input /home/flow/FlowData/userdata/bcounts/Project_snRNA Seq Trial_Skeletal Muscl_7/FeatureFilterResult-24943219446485782/filtered.mat.matrix --sample-ids /home/flow/FlowData/userdata/bcounts/Project_snRNA Seq Trial_Skeletal Muscl_7/FilterCellsResult-24942551988630052/filtered.observations --chart-output /home/flow/FlowData/userdata/bcounts/Project_snRNA Seq Trial_Skeletal Muscl_7/FeatureFilterResult-24943219446485782/expressionChartCache.json --stats-output /home/flow/FlowData/userdata/bcounts/Project_snRNA Seq Trial_Skeletal Muscl_7/FeatureFilterResult-24943219446485782/feature_distribution.txt --threads 4
java: setup
java: java
java: setup
java: java
java: if
java: setup
java: if
java: java

**PCA -> Graph-based clustering -> compute biomarkers**
java: setup
java: setup
/opt/partek_flow/bin/gene_analysis/prepare_feature_rank --parameters /home/flow/FlowData/userdata/bcounts/Project_snRNA Seq Trial_Skeletal Muscl_7/FeatureRankResult-24947351935946188/parameters15167242062087776248.json --attribute Graph-based --contrast 1 --output /home/flow/FlowData/userdata/bcounts/Project_snRNA Seq Trial_Skeletal Muscl_7/FeatureRankResult-24947351935946188/attribute11290147971621120944.txt

/opt/partek_flow/bin/gene_analysis/prepare_feature_rank --parameters /home/flow/FlowData/userdata/bcounts/Project_snRNA Seq Trial_Skeletal Muscl_7/FeatureRankResult-24947351935946188/parameters15167242062087776248.json --attribute Graph-based --contrast 2 --output /home/flow/FlowData/userdata/bcounts/Project_snRNA Seq Trial_Skeletal Muscl_7/FeatureRankResult-24947351935946188/attribute374231948382260871.txt
/opt/partek_flow/bin/gene_analysis/prepare_feature_rank --parameters /home/flow/FlowData/userdata/bcounts/Project_snRNA Seq Trial_Skeletal Muscl_7/FeatureRankResult-24947351935946188/parameters15167242062087776248.json --attribute Graph-based --contrast 5 --output /home/flow/FlowData/userdata/bcounts/Project_snRNA Seq Trial_Skeletal Muscl_7/FeatureRankResult-24947351935946188/attribute9117197038414022559.txt
/opt/partek_flow/bin/gene_analysis/prepare_feature_rank --parameters /home/flow/FlowData/userdata/bcounts/Project_snRNA Seq Trial_Skeletal Muscl_7/FeatureRankResult-24947351935946188/parameters15167242062087776248.json --attribute Graph-based --contrast 10 --output /home/flow/FlowData/userdata/bcounts/Project_snRNA Seq Trial_Skeletal Muscl_7/FeatureRankResult-24947351935946188/attribute7800971883106173934.txt
/opt/partek_flow/bin/gene_analysis/prepare_feature_rank --parameters /home/flow/FlowData/userdata/bcounts/Project_snRNA Seq Trial_Skeletal Muscl_7/FeatureRankResult-24947351935946188/parameters15167242062087776248.json --attribute Graph-based --contrast 8 --output /home/flow/FlowData/userdata/bcounts/Project_snRNA Seq Trial_Skeletal Muscl_7/FeatureRankResult-24947351935946188/attribute5452947118439529055.txt
/opt/partek_flow/bin/gene_analysis/prepare_feature_rank --parameters /home/flow/FlowData/userdata/bcounts/Project_snRNA Seq Trial_Skeletal Muscl_7/FeatureRankResult-24947351935946188/parameters15167242062087776248.json --attribute Graph-based --contrast 3 --output /home/flow/FlowData/userdata/bcounts/Project_snRNA Seq Trial_Skeletal Muscl_7/FeatureRankResult-24947351935946188/attribute13785646312068876877.txt
/opt/partek_flow/bin/gene_analysis/prepare_feature_rank --parameters /home/flow/FlowData/userdata/bcounts/Project_snRNA Seq Trial_Skeletal Muscl_7/FeatureRankResult-24947351935946188/parameters15167242062087776248.json --attribute Graph-based --contrast 12 --output /home/flow/FlowData/userdata/bcounts/Project_snRNA Seq Trial_Skeletal Muscl_7/FeatureRankResult-24947351935946188/attribute14894001159712628492.txt
/opt/partek_flow/bin/gene_analysis/prepare_feature_rank --parameters /home/flow/FlowData/userdata/bcounts/Project_snRNA Seq Trial_Skeletal Muscl_7/FeatureRankResult-24947351935946188/parameters15167242062087776248.json --attribute Graph-based --contrast 11 --output /home/flow/FlowData/userdata/bcounts/Project_snRNA Seq Trial_Skeletal Muscl_7/FeatureRankResult-24947351935946188/attribute9122851081632224428.txt
/opt/partek_flow/bin/gene_analysis/prepare_feature_rank --parameters /home/flow/FlowData/userdata/bcounts/Project_snRNA Seq Trial_Skeletal Muscl_7/FeatureRankResult-24947351935946188/parameters15167242062087776248.json --attribute Graph-based --contrast 9 --output /home/flow/FlowData/userdata/bcounts/Project_snRNA Seq Trial_Skeletal Muscl_7/FeatureRankResult-24947351935946188/attribute9159158860954487680.txt
/opt/partek_flow/bin/gene_analysis/prepare_feature_rank --parameters /home/flow/FlowData/userdata/bcounts/Project_snRNA Seq Trial_Skeletal Muscl_7/FeatureRankResult-24947351935946188/parameters15167242062087776248.json --attribute Graph-based --contrast 4 --output /home/flow/FlowData/userdata/bcounts/Project_snRNA Seq Trial_Skeletal Muscl_7/FeatureRankResult-24947351935946188/attribute11129900165727528918.txt
/opt/partek_flow/bin/gene_analysis/prepare_feature_rank --parameters /home/flow/FlowData/userdata/bcounts/Project_snRNA Seq Trial_Skeletal Muscl_7/FeatureRankResult-24947351935946188/parameters15167242062087776248.json --attribute Graph-based --contrast 13 --output /home/flow/FlowData/userdata/bcounts/Project_snRNA Seq Trial_Skeletal Muscl_7/FeatureRankResult-24947351935946188/attribute16966506272795976789.txt
/opt/partek_flow/bin/gene_analysis/prepare_feature_rank --parameters /home/flow/FlowData/userdata/bcounts/Project_snRNA Seq Trial_Skeletal Muscl_7/FeatureRankResult-24947351935946188/parameters15167242062087776248.json --attribute Graph-based --contrast 7 --output /home/flow/FlowData/userdata/bcounts/Project_snRNA Seq Trial_Skeletal Muscl_7/FeatureRankResult-24947351935946188/attribute11772404276072744012.txt
/opt/partek_flow/bin/gene_analysis/prepare_feature_rank --parameters /home/flow/FlowData/userdata/bcounts/Project_snRNA Seq Trial_Skeletal Muscl_7/FeatureRankResult-24947351935946188/parameters15167242062087776248.json --attribute Graph-based --contrast 6 --output /home/flow/FlowData/userdata/bcounts/Project_snRNA Seq Trial_Skeletal Muscl_7/FeatureRankResult-24947351935946188/attribute7245712505701557244.txt
java: java
java: java
java: java
java: java
java: java
java: java
java: java
/opt/partek_flow/bin/gene_analysis/flow_diff_expression --input_bdb /home/flow/FlowData/userdata/bcounts/Project_snRNA Seq Trial_Skeletal Muscl_7/FeatureRankResult-24947351935946188/input14480121302757761393.json --model_config /home/flow/FlowData/userdata/bcounts/Project_snRNA Seq Trial_Skeletal Muscl_7/FeatureRankResult-24947351935946188/modelsComparisons1523297314452908925.json --attributes /home/flow/FlowData/userdata/bcounts/Project_snRNA Seq Trial_Skeletal Muscl_7/FeatureRankResult-24947351935946188/parameters11984485824489301643.json --output_folder

/home/flow/FlowData/userdata/bcounts/Project_snRNA Seq Trial_Skeletal Muscl_7/FeatureRankResult-24947351935946188/out550682547800171625 --df 0 -c AICc --lognormal --useFstat -l gene-level --logbase 2.0 --threads 0
/opt/partek_flow/bin/gene_analysis/flow_diff_expression --input_bdb /home/flow/FlowData/userdata/bcounts/Project_snRNA Seq Trial_Skeletal Muscl_7/FeatureRankResult-24947351935946188/input14480121302757761393.json --model_config /home/flow/FlowData/userdata/bcounts/Project_snRNA Seq Trial_Skeletal Muscl_7/FeatureRankResult-24947351935946188/modelsComparisons1523297314452908925.json --attributes /home/flow/FlowData/userdata/bcounts/Project_snRNA Seq Trial_Skeletal Muscl_7/FeatureRankResult-24947351935946188/parameters3788802976319220684.json --output_folder /home/flow/FlowData/userdata/bcounts/Project_snRNA Seq Trial_Skeletal Muscl_7/FeatureRankResult-24947351935946188/out12755245251948666718 --df 0 -c AICc --lognormal --useFstat -l gene-level --logbase 2.0 --threads 0
/opt/partek_flow/bin/gene_analysis/flow_diff_expression --input_bdb /home/flow/FlowData/userdata/bcounts/Project_snRNA Seq Trial_Skeletal Muscl_7/FeatureRankResult-24947351935946188/input14480121302757761393.json --model_config /home/flow/FlowData/userdata/bcounts/Project_snRNA Seq Trial_Skeletal Muscl_7/FeatureRankResult-24947351935946188/modelsComparisons1523297314452908925.json --attributes /home/flow/FlowData/userdata/bcounts/Project_snRNA Seq Trial_Skeletal Muscl_7/FeatureRankResult-24947351935946188/parameters10751672045123106544.json --output_folder /home/flow/FlowData/userdata/bcounts/Project_snRNA Seq Trial_Skeletal Muscl_7/FeatureRankResult-24947351935946188/out3096255215773938411 --df 0 -c AICc --lognormal --useFstat -l gene-level --logbase 2.0 --threads 0
/opt/partek_flow/bin/gene_analysis/flow_diff_expression --input_bdb /home/flow/FlowData/userdata/bcounts/Project_snRNA Seq Trial_Skeletal Muscl_7/FeatureRankResult-24947351935946188/input14480121302757761393.json --model_config /home/flow/FlowData/userdata/bcounts/Project_snRNA Seq Trial_Skeletal Muscl_7/FeatureRankResult-24947351935946188/modelsComparisons1523297314452908925.json --attributes /home/flow/FlowData/userdata/bcounts/Project_snRNA Seq Trial_Skeletal Muscl_7/FeatureRankResult-24947351935946188/parameters14946232440674347133.json --output_folder /home/flow/FlowData/userdata/bcounts/Project_snRNA Seq Trial_Skeletal Muscl_7/FeatureRankResult-24947351935946188/out6653479661713454191 --df 0 -c AICc --lognormal --useFstat -l gene-level --logbase 2.0 --threads 0
/opt/partek_flow/bin/gene_analysis/flow_diff_expression --input_bdb /home/flow/FlowData/userdata/bcounts/Project_snRNA Seq Trial_Skeletal Muscl_7/FeatureRankResult-24947351935946188/input14480121302757761393.json --model_config /home/flow/FlowData/userdata/bcounts/Project_snRNA Seq Trial_Skeletal Muscl_7/FeatureRankResult-24947351935946188/modelsComparisons1523297314452908925.json --attributes /home/flow/FlowData/userdata/bcounts/Project_snRNA Seq Trial_Skeletal Muscl_7/FeatureRankResult-24947351935946188/parameters14240539518245081184.json --output_folder /home/flow/FlowData/userdata/bcounts/Project_snRNA Seq Trial_Skeletal Muscl_7/FeatureRankResult-24947351935946188/out14312407273858412766 --df 0 -c AICc --lognormal --useFstat -l gene-level --logbase 2.0 --threads 0
/opt/partek_flow/bin/gene_analysis/flow_diff_expression --input_bdb /home/flow/FlowData/userdata/bcounts/Project_snRNA Seq Trial_Skeletal Muscl_7/FeatureRankResult-24947351935946188/input14480121302757761393.json --model_config /home/flow/FlowData/userdata/bcounts/Project_snRNA Seq Trial_Skeletal Muscl_7/FeatureRankResult-24947351935946188/modelsComparisons1523297314452908925.json --attributes /home/flow/FlowData/userdata/bcounts/Project_snRNA Seq Trial_Skeletal Muscl_7/FeatureRankResult-24947351935946188/parameters8899742831170473596.json --output_folder /home/flow/FlowData/userdata/bcounts/Project_snRNA Seq Trial_Skeletal Muscl_7/FeatureRankResult-24947351935946188/out11342194214335754876 --df 0 -c AICc --lognormal --useFstat -l gene-level --logbase 2.0 --threads 0
/opt/partek_flow/bin/gene_analysis/flow_diff_expression --input_bdb /home/flow/FlowData/userdata/bcounts/Project_snRNA Seq Trial_Skeletal Muscl_7/FeatureRankResult-24947351935946188/input14480121302757761393.json --model_config /home/flow/FlowData/userdata/bcounts/Project_snRNA Seq Trial_Skeletal Muscl_7/FeatureRankResult-24947351935946188/modelsComparisons1523297314452908925.json --attributes /home/flow/FlowData/userdata/bcounts/Project_snRNA Seq Trial_Skeletal Muscl_7/FeatureRankResult-24947351935946188/parameters7850906279972579753.json --output_folder /home/flow/FlowData/userdata/bcounts/Project_snRNA Seq Trial_Skeletal Muscl_7/FeatureRankResult-24947351935946188/out18124456167954724506 --df 0 -c AICc --lognormal --useFstat -l gene-level --logbase 2.0 --threads 0
java: java
java: java
java: java
java: java
java: java
/opt/partek_flow/bin/gene_analysis/flow_diff_expression --input_bdb /home/flow/FlowData/userdata/bcounts/Project_snRNA Seq Trial_Skeletal Muscl_7/FeatureRankResult-24947351935946188/input14480121302757761393.json --model_config /home/flow/FlowData/userdata/bcounts/Project_snRNA Seq Trial_Skeletal Muscl_7/FeatureRankResult-24947351935946188/modelsComparisons1523297314452908925.json --attributes /home/flow/FlowData/userdata/bcounts/Project_snRNA Seq Trial_Skeletal Muscl_7/FeatureRankResult-24947351935946188/parameters15069661436000754020.json --output_folder

/home/flow/FlowData/userdata/bcounts/Project_snRNA Seq Trial_Skeletal Muscl_7/FeatureRankResult-24947351935946188/out9053577097021944646 --df 0 -c AICc --lognormal --useFstat -l gene-level --logbase 2.0 --threads 0
/opt/partek_flow/bin/gene_analysis/flow_diff_expression --input_bdb /home/flow/FlowData/userdata/bcounts/Project_snRNA Seq Trial_Skeletal Muscl_7/FeatureRankResult-24947351935946188/input14480121302757761393.json --model_config /home/flow/FlowData/userdata/bcounts/Project_snRNA Seq Trial_Skeletal Muscl_7/FeatureRankResult-24947351935946188/modelsComparisons1523297314452908925.json --attributes /home/flow/FlowData/userdata/bcounts/Project_snRNA Seq Trial_Skeletal Muscl_7/FeatureRankResult-24947351935946188/parameters6773869350934886088.json --output_folder /home/flow/FlowData/userdata/bcounts/Project_snRNA Seq Trial_Skeletal Muscl_7/FeatureRankResult-24947351935946188/out1047121502131496154 --df 0 -c AICc --lognormal --useFstat -l gene-level --logbase 2.0 --threads 0
/opt/partek_flow/bin/gene_analysis/flow_diff_expression --input_bdb /home/flow/FlowData/userdata/bcounts/Project_snRNA Seq Trial_Skeletal Muscl_7/FeatureRankResult-24947351935946188/input14480121302757761393.json --model_config /home/flow/FlowData/userdata/bcounts/Project_snRNA Seq Trial_Skeletal Muscl_7/FeatureRankResult-24947351935946188/modelsComparisons1523297314452908925.json --attributes /home/flow/FlowData/userdata/bcounts/Project_snRNA Seq Trial_Skeletal Muscl_7/FeatureRankResult-24947351935946188/parameters10209944727619967000.json --output_folder /home/flow/FlowData/userdata/bcounts/Project_snRNA Seq Trial_Skeletal Muscl_7/FeatureRankResult-24947351935946188/out10023262134829139049 --df 0 -c AICc --lognormal --useFstat -l gene-level --logbase 2.0 --threads 0
/opt/partek_flow/bin/gene_analysis/flow_diff_expression --input_bdb /home/flow/FlowData/userdata/bcounts/Project_snRNA Seq Trial_Skeletal Muscl_7/FeatureRankResult-24947351935946188/input14480121302757761393.json --model_config /home/flow/FlowData/userdata/bcounts/Project_snRNA Seq Trial_Skeletal Muscl_7/FeatureRankResult-24947351935946188/modelsComparisons1523297314452908925.json --attributes /home/flow/FlowData/userdata/bcounts/Project_snRNA Seq Trial_Skeletal Muscl_7/FeatureRankResult-24947351935946188/parameters5446443475557475189.json --output_folder /home/flow/FlowData/userdata/bcounts/Project_snRNA Seq Trial_Skeletal Muscl_7/FeatureRankResult-24947351935946188/out9564495862705438947 --df 0 -c AICc --lognormal --useFstat -l gene-level --logbase 2.0 --threads 0
/opt/partek_flow/bin/gene_analysis/flow_diff_expression --input_bdb /home/flow/FlowData/userdata/bcounts/Project_snRNA Seq Trial_Skeletal Muscl_7/FeatureRankResult-24947351935946188/input14480121302757761393.json --model_config /home/flow/FlowData/userdata/bcounts/Project_snRNA Seq Trial_Skeletal Muscl_7/FeatureRankResult-24947351935946188/modelsComparisons1523297314452908925.json --attributes /home/flow/FlowData/userdata/bcounts/Project_snRNA Seq Trial_Skeletal Muscl_7/FeatureRankResult-24947351935946188/parameters10905527066807961708.json --output_folder /home/flow/FlowData/userdata/bcounts/Project_snRNA Seq Trial_Skeletal Muscl_7/FeatureRankResult-24947351935946188/out5649581974731676895 --df 0 -c AICc --lognormal --useFstat -l gene-level --logbase 2.0 --threads 0
java: java
/opt/partek_flow/bin/gene_analysis/flow_diff_expression --input_bdb /home/flow/FlowData/userdata/bcounts/Project_snRNA Seq Trial_Skeletal Muscl_7/FeatureRankResult-24947351935946188/input14480121302757761393.json --model_config /home/flow/FlowData/userdata/bcounts/Project_snRNA Seq Trial_Skeletal Muscl_7/FeatureRankResult-24947351935946188/modelsComparisons1523297314452908925.json --attributes /home/flow/FlowData/userdata/bcounts/Project_snRNA Seq Trial_Skeletal Muscl_7/FeatureRankResult-24947351935946188/parameters7962750222907492724.json --output_folder /home/flow/FlowData/userdata/bcounts/Project_snRNA Seq Trial_Skeletal Muscl_7/FeatureRankResult-24947351935946188/out15694689559006437774 --df 0 -c AICc --lognormal --useFstat -l gene-level --logbase 2.0 --threads 0
java: java
java: java

**Myonuclei PCA -> Graph based clusters -> Computer biomarkers**
java: setup
/opt/partek_flow/bin/quantification/find_feature_indices --input /home/flow/FlowData/userdata/zimmers1/Project_BC_snRNA SKM_7/SplitByAttributeResult-60179866621774321/split.6.mat.matrix --output /home/flow/FlowData/userdata/zimmers1/Project_BC_snRNA SKM_7/FeatureRankResult-60264740086396148/columnFilter13274752177293799861.txt --threads 0
java: setup
/opt/partek_flow/bin/datablock/filter_datablock --input /home/flow/FlowData/userdata/zimmers1/Project_BC_snRNA SKM_7/FeatureRankResult-60264740086396148/input14122670748576103197.json --output /home/flow/FlowData/userdata/zimmers1/Project_BC_snRNA SKM_7/FeatureRankResult-60264740086396148/output12221969373204508689.json --column-filter /home/flow/FlowData/userdata/zimmers1/Project_BC_snRNA SKM_7/FeatureRankResult-60264740086396148/columnFilter13274752177293799861.txt --sort_indices --threads 0
java: java
java: else
java: if

java: setup
/opt/partek_flow/bin/gene_analysis/prepare_feature_rank --parameters /home/flow/FlowData/userdata/zimmers1/Project_BC_snRNA SKM_7/FeatureRankResult-60264740086396148/parameters14586637385305900313.json --specification /home/flow/FlowData/userdata/zimmers1/Project_BC_snRNA SKM_7/FeatureRankResult-60264740086396148/specification14753114101982144732.txt --output /home/flow/FlowData/userdata/zimmers1/Project_BC_snRNA SKM_7/FeatureRankResult-60264740086396148/attribute7078700555750782970.txt
/opt/partek_flow/bin/gene_analysis/prepare_feature_rank --parameters /home/flow/FlowData/userdata/zimmers1/Project_BC_snRNA SKM_7/FeatureRankResult-60264740086396148/parameters14586637385305900313.json --specification /home/flow/FlowData/userdata/zimmers1/Project_BC_snRNA SKM_7/FeatureRankResult-60264740086396148/specification9619875534834328776.txt --output /home/flow/FlowData/userdata/zimmers1/Project_BC_snRNA SKM_7/FeatureRankResult-60264740086396148/attribute13222347986941906849.txt
java: java
java: java
/opt/partek_flow/bin/gene_analysis/flow_diff_expression --input_bdb /home/flow/FlowData/userdata/zimmers1/Project_BC_snRNA SKM_7/FeatureRankResult-60264740086396148/input17125353169713634807.json --model_config /home/flow/FlowData/userdata/zimmers1/Project_BC_snRNA SKM_7/FeatureRankResult-60264740086396148/modelsComparisons3841463429050753565.json --attributes /home/flow/FlowData/userdata/zimmers1/Project_BC_snRNA SKM_7/FeatureRankResult-60264740086396148/parameters11939941638833145035.json --output_folder /home/flow/FlowData/userdata/zimmers1/Project_BC_snRNA SKM_7/FeatureRankResult-60264740086396148/out10436155851898010484 -c AICc --lognormal --useFstat --logbase 2.0 --threads 0
/opt/partek_flow/bin/gene_analysis/prepare_feature_rank --parameters /home/flow/FlowData/userdata/zimmers1/Project_BC_snRNA SKM_7/FeatureRankResult-60264740086396148/parameters14586637385305900313.json --specification /home/flow/FlowData/userdata/zimmers1/Project_BC_snRNA SKM_7/FeatureRankResult-60264740086396148/specification3302759297112186527.txt --output /home/flow/FlowData/userdata/zimmers1/Project_BC_snRNA SKM_7/FeatureRankResult-60264740086396148/attribute14800582167227907671.txt
/opt/partek_flow/bin/gene_analysis/prepare_feature_rank --parameters /home/flow/FlowData/userdata/zimmers1/Project_BC_snRNA SKM_7/FeatureRankResult-60264740086396148/parameters14586637385305900313.json --specification /home/flow/FlowData/userdata/zimmers1/Project_BC_snRNA SKM_7/FeatureRankResult-60264740086396148/specification5497951397908770435.txt --output /home/flow/FlowData/userdata/zimmers1/Project_BC_snRNA SKM_7/FeatureRankResult-60264740086396148/attribute18140801039306164781.txt
/opt/partek_flow/bin/gene_analysis/prepare_feature_rank --parameters /home/flow/FlowData/userdata/zimmers1/Project_BC_snRNA SKM_7/FeatureRankResult-60264740086396148/parameters14586637385305900313.json --specification /home/flow/FlowData/userdata/zimmers1/Project_BC_snRNA SKM_7/FeatureRankResult-60264740086396148/specification15131598632399526478.txt --output /home/flow/FlowData/userdata/zimmers1/Project_BC_snRNA SKM_7/FeatureRankResult-60264740086396148/attribute11571556535664940747.txt
/opt/partek_flow/bin/gene_analysis/prepare_feature_rank --parameters /home/flow/FlowData/userdata/zimmers1/Project_BC_snRNA SKM_7/FeatureRankResult-60264740086396148/parameters14586637385305900313.json --specification /home/flow/FlowData/userdata/zimmers1/Project_BC_snRNA SKM_7/FeatureRankResult-60264740086396148/specification1534271916873353056.txt --output /home/flow/FlowData/userdata/zimmers1/Project_BC_snRNA SKM_7/FeatureRankResult-60264740086396148/attribute15729806628245664432.txt
/opt/partek_flow/bin/gene_analysis/prepare_feature_rank --parameters /home/flow/FlowData/userdata/zimmers1/Project_BC_snRNA SKM_7/FeatureRankResult-60264740086396148/parameters14586637385305900313.json --specification /home/flow/FlowData/userdata/zimmers1/Project_BC_snRNA SKM_7/FeatureRankResult-60264740086396148/specification5236950474898030800.txt --output /home/flow/FlowData/userdata/zimmers1/Project_BC_snRNA SKM_7/FeatureRankResult-60264740086396148/attribute17843493462274969460.txt
/opt/partek_flow/bin/gene_analysis/prepare_feature_rank --parameters /home/flow/FlowData/userdata/zimmers1/Project_BC_snRNA SKM_7/FeatureRankResult-60264740086396148/parameters14586637385305900313.json --specification /home/flow/FlowData/userdata/zimmers1/Project_BC_snRNA SKM_7/FeatureRankResult-60264740086396148/specification11192398871920815578.txt --output /home/flow/FlowData/userdata/zimmers1/Project_BC_snRNA SKM_7/FeatureRankResult-60264740086396148/attribute730039509749509694.txt
/opt/partek_flow/bin/gene_analysis/flow_diff_expression --input_bdb /home/flow/FlowData/userdata/zimmers1/Project_BC_snRNA SKM_7/FeatureRankResult-60264740086396148/input17125353169713634807.json --model_config /home/flow/FlowData/userdata/zimmers1/Project_BC_snRNA SKM_7/FeatureRankResult-60264740086396148/modelsComparisons3841463429050753565.json --attributes /home/flow/FlowData/userdata/zimmers1/Project_BC_snRNA SKM_7/FeatureRankResult-60264740086396148/parameters12261668855342612644.json --output_folder /home/flow/FlowData/userdata/zimmers1/Project_BC_snRNA SKM_7/FeatureRankResult-60264740086396148/out2839689076811421235 -c AICc --lognormal --useFstat --logbase 2.0 --threads 0
java: java
/opt/partek_flow/bin/gene_analysis/flow_diff_expression --input_bdb /home/flow/FlowData/userdata/zimmers1/Project_BC_snRNA SKM_7/FeatureRankResult-60264740086396148/input17125353169713634807.json --model_config

/home/flow/FlowData/userdata/zimmers1/Project_BC_snRNA SKM_7/FeatureRankResult-60264740086396148/modelsComparisons3841463429050753565.json --attributes /home/flow/FlowData/userdata/zimmers1/Project_BC_snRNA SKM_7/FeatureRankResult-60264740086396148/parameters10934149107715311039.json --output_folder /home/flow/FlowData/userdata/zimmers1/Project_BC_snRNA SKM_7/FeatureRankResult-60264740086396148/out482624087476794906 -c AICc --lognormal --useFstat --logbase 2.0 --threads 0
java: java
/opt/partek_flow/bin/gene_analysis/flow_diff_expression --input_bdb /home/flow/FlowData/userdata/zimmers1/Project_BC_snRNA SKM_7/FeatureRankResult-60264740086396148/input17125353169713634807.json --model_config /home/flow/FlowData/userdata/zimmers1/Project_BC_snRNA SKM_7/FeatureRankResult-60264740086396148/modelsComparisons3841463429050753565.json --attributes /home/flow/FlowData/userdata/zimmers1/Project_BC_snRNA SKM_7/FeatureRankResult-60264740086396148/parameters8111636857144328389.json --output_folder /home/flow/FlowData/userdata/zimmers1/Project_BC_snRNA SKM_7/FeatureRankResult-60264740086396148/out9090124893741939098 -c AICc --lognormal --useFstat --logbase 2.0 --threads 0
java: java
java: java
java: java
/opt/partek_flow/bin/gene_analysis/flow_diff_expression --input_bdb /home/flow/FlowData/userdata/zimmers1/Project_BC_snRNA SKM_7/FeatureRankResult-60264740086396148/input17125353169713634807.json --model_config /home/flow/FlowData/userdata/zimmers1/Project_BC_snRNA SKM_7/FeatureRankResult-60264740086396148/modelsComparisons3841463429050753565.json --attributes /home/flow/FlowData/userdata/zimmers1/Project_BC_snRNA SKM_7/FeatureRankResult-60264740086396148/parameters9218916051036735379.json --output_folder /home/flow/FlowData/userdata/zimmers1/Project_BC_snRNA SKM_7/FeatureRankResult-60264740086396148/out2315388042318233122 -c AICc --lognormal --useFstat --logbase 2.0 --threads 0
/opt/partek_flow/bin/gene_analysis/flow_diff_expression --input_bdb /home/flow/FlowData/userdata/zimmers1/Project_BC_snRNA SKM_7/FeatureRankResult-60264740086396148/input17125353169713634807.json --model_config /home/flow/FlowData/userdata/zimmers1/Project_BC_snRNA SKM_7/FeatureRankResult-60264740086396148/modelsComparisons3841463429050753565.json --attributes /home/flow/FlowData/userdata/zimmers1/Project_BC_snRNA SKM_7/FeatureRankResult-60264740086396148/parameters11633483752045125789.json --output_folder /home/flow/FlowData/userdata/zimmers1/Project_BC_snRNA SKM_7/FeatureRankResult-60264740086396148/out1117178836410159949l -c AICc --lognormal --useFstat --logbase 2.0 --threads 0
java: java
/opt/partek_flow/bin/gene_analysis/flow_diff_expression --input_bdb /home/flow/FlowData/userdata/zimmers1/Project_BC_snRNA SKM_7/FeatureRankResult-60264740086396148/input17125353169713634807.json --model_config /home/flow/FlowData/userdata/zimmers1/Project_BC_snRNA SKM_7/FeatureRankResult-60264740086396148/modelsComparisons3841463429050753565.json --attributes /home/flow/FlowData/userdata/zimmers1/Project_BC_snRNA SKM_7/FeatureRankResult-60264740086396148/parameters5923204973747426692.json --output_folder /home/flow/FlowData/userdata/zimmers1/Project_BC_snRNA SKM_7/FeatureRankResult-60264740086396148/out9300263005556550322 -c AICc --lognormal --useFstat --logbase 2.0 --threads 0
/opt/partek_flow/bin/gene_analysis/flow_diff_expression --input_bdb /home/flow/FlowData/userdata/zimmers1/Project_BC_snRNA SKM_7/FeatureRankResult-60264740086396148/input17125353169713634807.json --model_config /home/flow/FlowData/userdata/zimmers1/Project_BC_snRNA SKM_7/FeatureRankResult-60264740086396148/modelsComparisons3841463429050753565.json --attributes /home/flow/FlowData/userdata/zimmers1/Project_BC_snRNA SKM_7/FeatureRankResult-60264740086396148/parameters13382814412108264239.json --output_folder /home/flow/FlowData/userdata/zimmers1/Project_BC_snRNA SKM_7/FeatureRankResult-60264740086396148/out5087920853911466802 -c AICc --lognormal --useFstat --logbase 2.0 --threads 0
java: java
java: java

**Human**
**Filter Cells**
java: setup
/opt/partek_flow/bin/tabular/flow_table --threads 0 -p /home/flow/FlowData/userdata/zimmers1/Project_SKMus73_Muscle snRNAseq_147/FilterCellsResult-2474541303352860/parameters13651264743228037558.json
java: setup
java: setup

/opt/partek_flow/bin/datablock/filter_datablock --input /home/flow/FlowData/userdata/zimmers1/Project_SKMus73_Muscle snRNAseq_147/FilterCellsResult-2474541303352860/input1824636517661630335.json --output /home/flow/FlowData/userdata/zimmers1/Project_SKMus73_Muscle snRNAseq_147/FilterCellsResult-2474541303352860/output16129465021989782674.json --row-vector-filter /home/flow/FlowData/userdata/zimmers1/Project_SKMus73_Muscle snRNAseq_147/FilterCellsResult-2474541303352860/rowIndices3833371787924024249.bin --sort_indices --threads 0
java: setup
java: java
java: setup
java: build_stats_command
java: setup
/opt/partek_flow/bin/datablock/datablock_stats --input /home/flow/FlowData/userdata/zimmers1/Project_SKMus73_Muscle snRNAseq_147/FilterCellsResult-2474541303352860/filtered.mat.matrix --sample-ids /home/flow/FlowData/userdata/zimmers1/Project_SKMus73_Muscle snRNAseq_147/FilterCellsResult-2474541303352860/filtered.observations --output /home/flow/FlowData/userdata/zimmers1/Project_SKMus73_Muscle snRNAseq_147/FilterCellsResult-2474541303352860/feature_distribution.txt --threads 0
java: java
java: java

/opt/partek_flow/bin/quantification/find_feature_indices --input /home/flow/FlowData/userdata/zimmers1/Project_SKMus73_Muscle snRNAseq_147/FilterCellsResult-2474541303352860/filtered.mat.matrix --output /home/flow/FlowData/userdata/zimmers1/Project_SKMus73_Muscle snRNAseq_147/FeatureFilterResult-2475120891405629/filter12580887428735767138.txt --threads 0 --expression_method value --operation le --expression_value 0.0 --expression_percent 99.0
java: setup
java: setup
/opt/partek_flow/bin/datablock/filter_datablock --input /home/flow/FlowData/userdata/zimmers1/Project_SKMus73_Muscle snRNAseq_147/FeatureFilterResult-2475120891405629/input16204797454175042826.json --output /home/flow/FlowData/userdata/zimmers1/Project_SKMus73_Muscle snRNAseq_147/FeatureFilterResult-2475120891405629/output8021556531113048527.json --column-filter /home/flow/FlowData/userdata/zimmers1/Project_SKMus73_Muscle snRNAseq_147/FeatureFilterResult-2475120891405629/filter12580887428735767138.txt --sort_indices --threads 0
java: if
java: setup
java: setup
java: setup
/opt/partek_flow/bin/single_cell/sc_qa_qc --input-data /home/flow/FlowData/userdata/zimmers1/Project_SKMus73_Muscle snRNAseq_147/FeatureFilterResult-2475120891405629/filtered.mat.matrix --input-features /home/flow/FlowData/userdata/zimmers1/Project_SKMus73_Muscle snRNAseq_147/FeatureFilterResult-2475120891405629/filtered.col.matrix --input-samples /home/flow/FlowData/userdata/zimmers1/Project_SKMus73_Muscle snRNAseq_147/FeatureFilterResult-2475120891405629/filtered.observations --input-sample-annotation /home/flow/FlowData/userdata/zimmers1/Project_SKMus73_Muscle snRNAseq_147/FeatureFilterResult-2475120891405629/filtered.row.annotation --compute-iff-exists --output /home/flow/FlowData/userdata/zimmers1/Project_SKMus73_Muscle snRNAseq_147/FeatureFilterResult-2475120891405629/scQAQC9355230995490730067.txt --threads 0
java: setup
/opt/partek_flow/bin/combine_datablocks/merge_attributes --threads 0 --specification /home/flow/FlowData/userdata/zimmers1/Project_SKMus73_Muscle snRNAseq_147/FeatureFilterResult-2475120891405629/Merge_Specification12723954041400978619.txt --input /home/flow/FlowData/userdata/zimmers1/Project_SKMus73_Muscle snRNAseq_147/FeatureFilterResult-2475120891405629/outputRowAnnotation982633641007461662.txt --output /home/flow/FlowData/userdata/zimmers1/Project_SKMus73_Muscle snRNAseq_147/FeatureFilterResult-2475120891405629/temp12062504860848241301.row.annotation
java: java
java: if
java: setup
/opt/partek_flow/bin/quantification/generate_expression_chart_data --input /home/flow/FlowData/userdata/zimmers1/Project_SKMus73_Muscle snRNAseq_147/FeatureFilterResult-2475120891405629/filtered.mat.matrix --sample-ids /home/flow/FlowData/userdata/zimmers1/Project_SKMus73_Muscle snRNAseq_147/FilterCellsResult-2474541303352860/filtered.observations --chart-output /home/flow/FlowData/userdata/zimmers1/Project_SKMus73_Muscle snRNAseq_147/FeatureFilterResult-

2475120891405629/expressionChartCache.json --stats-output /home/flow/FlowData/userdata/zimmers1/Project_SKMus73_Muscle snRNAseq_147/FeatureFilterResult-2475120891405629/feature_distribution.txt --threads 4
java: setup
java: java
java: setup
java: java
java: if
java: setup
java: build_stats_command
java: if
java: java

**Normalization**
/opt/partek_flow/bin/datablock/normalize_datablock --threads 0 --input /home/flow/FlowData/userdata/zimmers1/Project_SKMus73_Muscle snRNAseq_147/NormalizationResult-2475206368908850/input1094166287101618293l.json --output /home/flow/FlowData/userdata/zimmers1/Project_SKMus73_Muscle snRNAseq_147/NormalizationResult-2475206368908850/normalized.mat.matrix --transform-on Cells --total-count --add 1.0 --log 2.0 --output_effective_libsize /home/flow/FlowData/userdata/zimmers1/Project_SKMus73_Muscle snRNAseq_147/NormalizationResult-2475206368908850/normalized.effective_lib_sizes.txt
java: setup
java: java
java: setup
java: build_stats_command
java: if
/opt/partek_flow/bin/quantification/generate_expression_chart_data --input /home/flow/FlowData/userdata/zimmers1/Project_SKMus73_Muscle snRNAseq_147/NormalizationResult-2475206368908850/normalized.mat.matrix --sample-ids /home/flow/FlowData/userdata/zimmers1/Project_SKMus73_Muscle snRNAseq_147/FilterCellsResult-2474541303352860/filtered.observations --chart-output /home/flow/FlowData/userdata/zimmers1/Project_SKMus73_Muscle snRNAseq_147/NormalizationResult-2475206368908850/expressionChartCache.json --stats-output /home/flow/FlowData/userdata/zimmers1/Project_SKMus73_Muscle snRNAseq_147/NormalizationResult-2475206368908850/feature_distribution.txt --threads 4
java: setup
/opt/partek_flow/bin/quantification/generate_expression_chart_data --input /home/flow/FlowData/userdata/zimmers1/Project_SKMus73_Muscle snRNAseq_147/FeatureFilterResult-2475120891405629/filtered.mat.matrix --sample-ids /home/flow/FlowData/userdata/zimmers1/Project_SKMus73_Muscle snRNAseq_147/FilterCellsResult-2474541303352860/filtered.observations --chart-output /home/flow/FlowData/userdata/zimmers1/Project_SKMus73_Muscle snRNAseq_147/FeatureFilterResult-2475120891405629/expressionChartCache.json --threads 4
/opt/partek_flow/bin/datablock/datablock_stats --input /home/flow/FlowData/userdata/zimmers1/Project_SKMus73_Muscle snRNAseq_147/FeatureFilterResult-2475120891405629/filtered.mat.matrix --sample-ids /home/flow/FlowData/userdata/zimmers1/Project_SKMus73_Muscle snRNAseq_147/FilterCellsResult-2474541303352860/filtered.observations --output /home/flow/FlowData/userdata/zimmers1/Project_SKMus73_Muscle snRNAseq_147/NormalizationResult-2475206368908850/dataBlockStats6419507951170113221.txt --threads 0
java: build_stats_command
java: java

/opt/partek_flow/bin/quantification/find_feature_indices --input /home/flow/FlowData/userdata/zimmers1/Project_SKMus73_Muscle snRNAseq_147/NormalizationResult-2475206368908850/normalized.mat.matrix --output /home/flow/FlowData/project_output/Project_SKMus73_Muscle snRNAseq_147/PrincipalComponentAnalysisResult-2561726763331133/colFilter2366618650675815435.idx --stats_method variance --stats_value 2000 --threads 0
java: setup
/opt/partek_flow/bin/pca/quantPCA --number-pcs 100 --input-data /home/flow/FlowData/project_output/Project_SKMus73_Muscle snRNAseq_147/PrincipalComponentAnalysisResult-2561726763331133/input1165133056988l505081.json --col-filter /home/flow/FlowData/project_output/Project_SKMus73_Muscle snRNAseq_147/PrincipalComponentAnalysisResult-2561726763331133/colFilter2366618650675815435.idx --output-data /home/flow/FlowData/project_output/Project_SKMus73_Muscle snRNAseq_147/PrincipalComponentAnalysisResult-2561726763331133/output7485860717442310591.json --output-eigenvalues /home/flow/FlowData/project_output/Project_SKMus73_Muscle snRNAseq_147/PrincipalComponentAnalysisResult-2561726763331133/eigenvalues0.txt --output-loadings /home/flow/FlowData/project_output/Project_SKMus73_Muscle snRNAseq_147/PrincipalComponentAnalysisResult-2561726763331133/loadings0.txt --shift --threads 0
java: java
java: setup

java: setup
java: setup
java: setup
java: java
java: java
java: setup

**PCA -> Graph-based clustering -> compute biomarkers**
java: setup
/opt/partek_flow/bin/quantification/find_feature_indices --input /home/flow/FlowData/userdata/zimmers1/Project_SKMus73_Muscle snRNAseq_147/NormalizationResult-2475206368908850/normalized.mat.matrix --output /home/flow/FlowData/userdata/zimmers1/Project_SKMus73_Muscle snRNAseq_147/FeatureRankResult-2729041228720229/columnFilter10373758564869641512.txt --threads 0
java: setup
/opt/partek_flow/bin/datablock/filter_datablock --input /home/flow/FlowData/userdata/zimmers1/Project_SKMus73_Muscle snRNAseq_147/FeatureRankResult-2729041228720229/input14374985823550682551.json --output /home/flow/FlowData/userdata/zimmers1/Project_SKMus73_Muscle snRNAseq_147/FeatureRankResult-2729041228720229/output5348884955065691552.json --column-filter /home/flow/FlowData/userdata/zimmers1/Project_SKMus73_Muscle snRNAseq_147/FeatureRankResult-2729041228720229/columnFilter10373758564869641512.txt --sort_indices --threads 0
java: java
java: if
java: else
java: setup
/opt/partek_flow/bin/gene_analysis/prepare_feature_rank --parameters /home/flow/FlowData/userdata/zimmers1/Project_SKMus73_Muscle snRNAseq_147/FeatureRankResult-2729041228720229/parameters14774457014878304436.json --specification /home/flow/FlowData/userdata/zimmers1/Project_SKMus73_Muscle snRNAseq_147/FeatureRankResult-2729041228720229/specification12776646022467813345.txt --output /home/flow/FlowData/userdata/zimmers1/Project_SKMus73_Muscle snRNAseq_147/FeatureRankResult-2729041228720229/attribute3086564506794425559.txt
/opt/partek_flow/bin/gene_analysis/prepare_feature_rank --parameters /home/flow/FlowData/userdata/zimmers1/Project_SKMus73_Muscle snRNAseq_147/FeatureRankResult-2729041228720229/parameters14774457014878304436.json --specification /home/flow/FlowData/userdata/zimmers1/Project_SKMus73_Muscle snRNAseq_147/FeatureRankResult-2729041228720229/specification3134022398682308073.txt --output /home/flow/FlowData/userdata/zimmers1/Project_SKMus73_Muscle snRNAseq_147/FeatureRankResult-2729041228720229/attribute18174211095048151855.txt
/opt/partek_flow/bin/gene_analysis/prepare_feature_rank --parameters /home/flow/FlowData/userdata/zimmers1/Project_SKMus73_Muscle snRNAseq_147/FeatureRankResult-2729041228720229/parameters14774457014878304436.json --specification /home/flow/FlowData/userdata/zimmers1/Project_SKMus73_Muscle snRNAseq_147/FeatureRankResult-2729041228720229/specification17135628386002208273.txt --output /home/flow/FlowData/userdata/zimmers1/Project_SKMus73_Muscle snRNAseq_147/FeatureRankResult-2729041228720229/attribute18243556430131081408.txt
/opt/partek_flow/bin/gene_analysis/prepare_feature_rank --parameters /home/flow/FlowData/userdata/zimmers1/Project_SKMus73_Muscle snRNAseq_147/FeatureRankResult-2729041228720229/parameters14774457014878304436.json --specification /home/flow/FlowData/userdata/zimmers1/Project_SKMus73_Muscle snRNAseq_147/FeatureRankResult-2729041228720229/specification12837337360695451111.txt --output /home/flow/FlowData/userdata/zimmers1/Project_SKMus73_Muscle snRNAseq_147/FeatureRankResult-2729041228720229/attribute8021413064784668306.txt
/opt/partek_flow/bin/gene_analysis/prepare_feature_rank --parameters /home/flow/FlowData/userdata/zimmers1/Project_SKMus73_Muscle snRNAseq_147/FeatureRankResult-2729041228720229/parameters14774457014878304436.json --specification /home/flow/FlowData/userdata/zimmers1/Project_SKMus73_Muscle snRNAseq_147/FeatureRankResult-2729041228720229/specification18048340087250186542.txt --output /home/flow/FlowData/userdata/zimmers1/Project_SKMus73_Muscle snRNAseq_147/FeatureRankResult-2729041228720229/attribute9051911954989551206.txt
/opt/partek_flow/bin/gene_analysis/prepare_feature_rank --parameters /home/flow/FlowData/userdata/zimmers1/Project_SKMus73_Muscle snRNAseq_147/FeatureRankResult-2729041228720229/parameters14774457014878304436.json --specification

/home/flow/FlowData/userdata/zimmers1/Project_SKMus73_Muscle snRNAseq_147/FeatureRankResult-2729041228720229/specification15627992105712117652.txt --output
/home/flow/FlowData/userdata/zimmers1/Project_SKMus73_Muscle snRNAseq_147/FeatureRankResult-2729041228720229/attribute16354501834566200777.txt
/opt/partek_flow/bin/gene_analysis/prepare_feature_rank --parameters
/home/flow/FlowData/userdata/zimmers1/Project_SKMus73_Muscle snRNAseq_147/FeatureRankResult-2729041228720229/parameters14774457014878304436.json --specification
/home/flow/FlowData/userdata/zimmers1/Project_SKMus73_Muscle snRNAseq_147/FeatureRankResult-2729041228720229/specification7548254294473181738.txt --output
/home/flow/FlowData/userdata/zimmers1/Project_SKMus73_Muscle snRNAseq_147/FeatureRankResult-2729041228720229/attribute4947661653672327912.txt
/opt/partek_flow/bin/gene_analysis/prepare_feature_rank --parameters
/home/flow/FlowData/userdata/zimmers1/Project_SKMus73_Muscle snRNAseq_147/FeatureRankResult-2729041228720229/parameters14774457014878304436.json --specification
/home/flow/FlowData/userdata/zimmers1/Project_SKMus73_Muscle snRNAseq_147/FeatureRankResult-2729041228720229/specification9423729711397279035.txt --output
/home/flow/FlowData/userdata/zimmers1/Project_SKMus73_Muscle snRNAseq_147/FeatureRankResult-2729041228720229/attribute799867722934854764.txt
java: java
java: java
/opt/partek_flow/bin/gene_analysis/prepare_feature_rank --parameters
/home/flow/FlowData/userdata/zimmers1/Project_SKMus73_Muscle snRNAseq_147/FeatureRankResult-2729041228720229/parameters14774457014878304436.json --specification
/home/flow/FlowData/userdata/zimmers1/Project_SKMus73_Muscle snRNAseq_147/FeatureRankResult-2729041228720229/specification7252312437836122579.txt --output
/home/flow/FlowData/userdata/zimmers1/Project_SKMus73_Muscle snRNAseq_147/FeatureRankResult-2729041228720229/attribute12787785713445180661.txt
/opt/partek_flow/bin/gene_analysis/flow_diff_expression --input_bdb
/home/flow/FlowData/userdata/zimmers1/Project_SKMus73_Muscle snRNAseq_147/FeatureRankResult-2729041228720229/input7885337249785227464.json --model_config
/home/flow/FlowData/userdata/zimmers1/Project_SKMus73_Muscle snRNAseq_147/FeatureRankResult-2729041228720229/modelsComparisons11724411551003919310.json --attributes
/home/flow/FlowData/userdata/zimmers1/Project_SKMus73_Muscle snRNAseq_147/FeatureRankResult-2729041228720229/parameters8993891110016229924.json --output_folder
/home/flow/FlowData/userdata/zimmers1/Project_SKMus73_Muscle snRNAseq_147/FeatureRankResult-2729041228720229/out4359954288238045372 -c AICc --lognormal --useFstat --logbase 2.0 --threads 0
java: java
/opt/partek_flow/bin/gene_analysis/flow_diff_expression --input_bdb
/home/flow/FlowData/userdata/zimmers1/Project_SKMus73_Muscle snRNAseq_147/FeatureRankResult-2729041228720229/input7885337249785227464.json --model_config
/home/flow/FlowData/userdata/zimmers1/Project_SKMus73_Muscle snRNAseq_147/FeatureRankResult-2729041228720229/modelsComparisons11724411551003919310.json --attributes
/home/flow/FlowData/userdata/zimmers1/Project_SKMus73_Muscle snRNAseq_147/FeatureRankResult-2729041228720229/parameters3749122349680233017.json --output_folder
/home/flow/FlowData/userdata/zimmers1/Project_SKMus73_Muscle snRNAseq_147/FeatureRankResult-2729041228720229/out17690260730691549192 -c AICc --lognormal --useFstat --logbase 2.0 --threads 0
java: java
java: java
/opt/partek_flow/bin/gene_analysis/flow_diff_expression --input_bdb
/home/flow/FlowData/userdata/zimmers1/Project_SKMus73_Muscle snRNAseq_147/FeatureRankResult-2729041228720229/input7885337249785227464.json --model_config
/home/flow/FlowData/userdata/zimmers1/Project_SKMus73_Muscle snRNAseq_147/FeatureRankResult-2729041228720229/modelsComparisons11724411551003919310.json --attributes
/home/flow/FlowData/userdata/zimmers1/Project_SKMus73_Muscle snRNAseq_147/FeatureRankResult-2729041228720229/parameters12058895408013998303.json --output_folder
/home/flow/FlowData/userdata/zimmers1/Project_SKMus73_Muscle snRNAseq_147/FeatureRankResult-2729041228720229/out2282420960812510733 -c AICc --lognormal --useFstat --logbase 2.0 --threads 0
/opt/partek_flow/bin/gene_analysis/flow_diff_expression --input_bdb
/home/flow/FlowData/userdata/zimmers1/Project_SKMus73_Muscle snRNAseq_147/FeatureRankResult-2729041228720229/input7885337249785227464.json --model_config
/home/flow/FlowData/userdata/zimmers1/Project_SKMus73_Muscle snRNAseq_147/FeatureRankResult-2729041228720229/modelsComparisons11724411551003919310.json --attributes

/home/flow/FlowData/userdata/zimmers1/Project_SKMus73_Muscle snRNAseq_147/FeatureRankResult-2729041228720229/parameters1783293521857765O484.json --output_folder
/home/flow/FlowData/userdata/zimmers1/Project_SKMus73_Muscle snRNAseq_147/FeatureRankResult-2729041228720229/out1593972875O591873054 -c AICc --lognormal --useFstat --logbase 2.0 --threads 0
java: java
/opt/partek_flow/bin/gene_analysis/flow_diff_expression --input_bdb
/home/flow/FlowData/userdata/zimmers1/Project_SKMus73_Muscle snRNAseq_147/FeatureRankResult-2729041228720229/input7885337249785227464.json --model_config
/home/flow/FlowData/userdata/zimmers1/Project_SKMus73_Muscle snRNAseq_147/FeatureRankResult-2729041228720229/modelsComparisons1172441155100391931O.json --attributes
/home/flow/FlowData/userdata/zimmers1/Project_SKMus73_Muscle snRNAseq_147/FeatureRankResult-2729041228720229/parameters1391257721751O330837.json --output_folder
/home/flow/FlowData/userdata/zimmers1/Project_SKMus73_Muscle snRNAseq_147/FeatureRankResult-2729041228720229/out8253413384917405650 -c AICc --lognormal --useFstat --logbase 2.0 --threads 0
java: java
/opt/partek_flow/bin/gene_analysis/flow_diff_expression --input_bdb
/home/flow/FlowData/userdata/zimmers1/Project_SKMus73_Muscle snRNAseq_147/FeatureRankResult-2729041228720229/input7885337249785227464.json --model_config
/home/flow/FlowData/userdata/zimmers1/Project_SKMus73_Muscle snRNAseq_147/FeatureRankResult-2729041228720229/modelsComparisons1172441155100391931O.json --attributes
/home/flow/FlowData/userdata/zimmers1/Project_SKMus73_Muscle snRNAseq_147/FeatureRankResult-2729041228720229/parameters8215639171092794434.json --output_folder
/home/flow/FlowData/userdata/zimmers1/Project_SKMus73_Muscle snRNAseq_147/FeatureRankResult-2729041228720229/out1547695568313982861 6 -c AICc --lognormal --useFstat --logbase 2.0 --threads 0
java: java
/opt/partek_flow/bin/gene_analysis/prepare_feature_rank --parameters
/home/flow/FlowData/userdata/zimmers1/Project_SKMus73_Muscle snRNAseq_147/FeatureRankResult-2729041228720229/parameters14774457014878304436.json --specification
/home/flow/FlowData/userdata/zimmers1/Project_SKMus73_Muscle snRNAseq_147/FeatureRankResult-2729041228720229/specification1727996593431570438 5.txt --output
/home/flow/FlowData/userdata/zimmers1/Project_SKMus73_Muscle snRNAseq_147/FeatureRankResult-2729041228720229/attribute12280780969392431420.txt
java: java
java: java
/opt/partek_flow/bin/gene_analysis/flow_diff_expression --input_bdb
/home/flow/FlowData/userdata/zimmers1/Project_SKMus73_Muscle snRNAseq_147/FeatureRankResult-2729041228720229/input7885337249785227464.json --model_config
/home/flow/FlowData/userdata/zimmers1/Project_SKMus73_Muscle snRNAseq_147/FeatureRankResult-2729041228720229/modelsComparisons1172441155100391931O.json --attributes
/home/flow/FlowData/userdata/zimmers1/Project_SKMus73_Muscle snRNAseq_147/FeatureRankResult-2729041228720229/parameters10933729868098907659.json --output_folder
/home/flow/FlowData/userdata/zimmers1/Project_SKMus73_Muscle snRNAseq_147/FeatureRankResult-2729041228720229/out11515970780308946845 -c AICc --lognormal --useFstat --logbase 2.0 --threads 0
/opt/partek_flow/bin/gene_analysis/flow_diff_expression --input_bdb
/home/flow/FlowData/userdata/zimmers1/Project_SKMus73_Muscle snRNAseq_147/FeatureRankResult-2729041228720229/input7885337249785227464.json --model_config
/home/flow/FlowData/userdata/zimmers1/Project_SKMus73_Muscle snRNAseq_147/FeatureRankResult-2729041228720229/modelsComparisons1172441155100391931O.json --attributes
/home/flow/FlowData/userdata/zimmers1/Project_SKMus73_Muscle snRNAseq_147/FeatureRankResult-2729041228720229/parameters484294335474433393.json --output_folder
/home/flow/FlowData/userdata/zimmers1/Project_SKMus73_Muscle snRNAseq_147/FeatureRankResult-2729041228720229/out4746010374249433536 -c AICc --lognormal --useFstat --logbase 2.0 --threads 0
/opt/partek_flow/bin/gene_analysis/flow_diff_expression --input_bdb
/home/flow/FlowData/userdata/zimmers1/Project_SKMus73_Muscle snRNAseq_147/FeatureRankResult-2729041228720229/input7885337249785227464.json --model_config
/home/flow/FlowData/userdata/zimmers1/Project_SKMus73_Muscle snRNAseq_147/FeatureRankResult-2729041228720229/modelsComparisons1172441155100391931O.json --attributes
/home/flow/FlowData/userdata/zimmers1/Project_SKMus73_Muscle snRNAseq_147/FeatureRankResult-2729041228720229/parameters9204843525575971048.json --output_folder
/home/flow/FlowData/userdata/zimmers1/Project_SKMus73_Muscle snRNAseq_147/FeatureRankResult-2729041228720229/out4162187687169730013 -c AICc --lognormal --useFstat --logbase 2.0 --threads 0

/opt/partek_flow/bin/gene_analysis/flow_diff_expression --input_bdb
/home/flow/FlowData/userdata/zimmers1/Project_SKMus73_Muscle snRNAseq_147/FeatureRankResult-2729041228720229/input7885337249785227464.json --model_config
/home/flow/FlowData/userdata/zimmers1/Project_SKMus73_Muscle snRNAseq_147/FeatureRankResult-2729041228720229/modelsComparisons11724411551003919310.json --attributes
/home/flow/FlowData/userdata/zimmers1/Project_SKMus73_Muscle snRNAseq_147/FeatureRankResult-2729041228720229/parameters2477946367923090445.json --output_folder
/home/flow/FlowData/userdata/zimmers1/Project_SKMus73_Muscle snRNAseq_147/FeatureRankResult-2729041228720229/out3664529342432564395 -c AICc --lognormal --useFstat --logbase 2.0 --threads 0
java: java
java: java

**Myonuclei PCA -> Graph-based clustering -> compute biomarkers**
java: setup
/opt/partek_flow/bin/quantification/find_feature_indices --input /home/flow/FlowData/userdata/zimmers1/Project_SKMus73_Muscle snRNAseq_147/SplitByAttributeResult-2733298647949985/split.4.mat.matrix --output
/home/flow/FlowData/userdata/zimmers1/Project_SKMus73_Muscle snRNAseq_147/FeatureRankResult-2734589615408625/columnFilter1608652827545515314.txt --threads 0
java: setup
/opt/partek_flow/bin/datablock/filter_datablock --input /home/flow/FlowData/userdata/zimmers1/Project_SKMus73_Muscle snRNAseq_147/FeatureRankResult-2734589615408625/input14872207109946404319.json --output
/home/flow/FlowData/userdata/zimmers1/Project_SKMus73_Muscle snRNAseq_147/FeatureRankResult-2734589615408625/output7057182428608404417.json --column-filter
/home/flow/FlowData/userdata/zimmers1/Project_SKMus73_Muscle snRNAseq_147/FeatureRankResult-2734589615408625/columnFilter1608652827545515314.txt --sort_indices --threads 0
java: java
java: if
java: else
java: setup
/opt/partek_flow/bin/gene_analysis/prepare_feature_rank --parameters
/home/flow/FlowData/userdata/zimmers1/Project_SKMus73_Muscle snRNAseq_147/FeatureRankResult-2734589615408625/parameters9275683716048498017.json --specification
/home/flow/FlowData/userdata/zimmers1/Project_SKMus73_Muscle snRNAseq_147/FeatureRankResult-2734589615408625/specification3791459420853817062.txt --output
/home/flow/FlowData/userdata/zimmers1/Project_SKMus73_Muscle snRNAseq_147/FeatureRankResult-2734589615408625/attribute5561166618590942131.txt
/opt/partek_flow/bin/gene_analysis/prepare_feature_rank --parameters
/home/flow/FlowData/userdata/zimmers1/Project_SKMus73_Muscle snRNAseq_147/FeatureRankResult-2734589615408625/parameters9275683716048498017.json --specification
/home/flow/FlowData/userdata/zimmers1/Project_SKMus73_Muscle snRNAseq_147/FeatureRankResult-2734589615408625/specification14408886663016648788.txt --output
/home/flow/FlowData/userdata/zimmers1/Project_SKMus73_Muscle snRNAseq_147/FeatureRankResult-2734589615408625/attribute7600386534878352208.txt
java: java
/opt/partek_flow/bin/gene_analysis/flow_diff_expression --input_bdb
/home/flow/FlowData/userdata/zimmers1/Project_SKMus73_Muscle snRNAseq_147/FeatureRankResult-2734589615408625/input3690637414517808576.json --model_config
/home/flow/FlowData/userdata/zimmers1/Project_SKMus73_Muscle snRNAseq_147/FeatureRankResult-2734589615408625/modelsComparisons11709814908268713496.json --attributes
/home/flow/FlowData/userdata/zimmers1/Project_SKMus73_Muscle snRNAseq_147/FeatureRankResult-2734589615408625/parameters4854012518532939662.json --output_folder
/home/flow/FlowData/userdata/zimmers1/Project_SKMus73_Muscle snRNAseq_147/FeatureRankResult-2734589615408625/out9124998137237480275 -c AICc --lognormal --useFstat --logbase 2.0 --threads 0
/opt/partek_flow/bin/gene_analysis/prepare_feature_rank --parameters
/home/flow/FlowData/userdata/zimmers1/Project_SKMus73_Muscle snRNAseq_147/FeatureRankResult-2734589615408625/parameters9275683716048498017.json --specification
/home/flow/FlowData/userdata/zimmers1/Project_SKMus73_Muscle snRNAseq_147/FeatureRankResult-2734589615408625/specification7554342128377432454.txt --output
/home/flow/FlowData/userdata/zimmers1/Project_SKMus73_Muscle snRNAseq_147/FeatureRankResult-2734589615408625/attribute14898327174954456913.txt

/opt/partek_flow/bin/gene_analysis/prepare_feature_rank --parameters
/home/flow/FlowData/userdata/zimmers1/Project_SKMus73_Muscle snRNAseq_147/FeatureRankResult-2734589615408625/parameters9275683716048498017.json --specification
/home/flow/FlowData/userdata/zimmers1/Project_SKMus73_Muscle snRNAseq_147/FeatureRankResult-2734589615408625/specification10143008467492924401.txt --output
/home/flow/FlowData/userdata/zimmers1/Project_SKMus73_Muscle snRNAseq_147/FeatureRankResult-2734589615408625/attribute6670197070479221504.txt
/opt/partek_flow/bin/gene_analysis/prepare_feature_rank --parameters
/home/flow/FlowData/userdata/zimmers1/Project_SKMus73_Muscle snRNAseq_147/FeatureRankResult-2734589615408625/parameters9275683716048498017.json --specification
/home/flow/FlowData/userdata/zimmers1/Project_SKMus73_Muscle snRNAseq_147/FeatureRankResult-2734589615408625/specification4357193358186031048.txt --output
/home/flow/FlowData/userdata/zimmers1/Project_SKMus73_Muscle snRNAseq_147/FeatureRankResult-2734589615408625/attribute1532353444403822034.txt
java: java
/opt/partek_flow/bin/gene_analysis/prepare_feature_rank --parameters
/home/flow/FlowData/userdata/zimmers1/Project_SKMus73_Muscle snRNAseq_147/FeatureRankResult-2734589615408625/parameters9275683716048498017.json --specification
/home/flow/FlowData/userdata/zimmers1/Project_SKMus73_Muscle snRNAseq_147/FeatureRankResult-2734589615408625/specification3759965807244051581.txt --output
/home/flow/FlowData/userdata/zimmers1/Project_SKMus73_Muscle snRNAseq_147/FeatureRankResult-2734589615408625/attribute17324101009601217789.txt
/opt/partek_flow/bin/gene_analysis/flow_diff_expression --input_bdb
/home/flow/FlowData/userdata/zimmers1/Project_SKMus73_Muscle snRNAseq_147/FeatureRankResult-2734589615408625/input3690637414517808576.json --model_config
/home/flow/FlowData/userdata/zimmers1/Project_SKMus73_Muscle snRNAseq_147/FeatureRankResult-2734589615408625/modelsComparisons11709814908268713496.json --attributes
/home/flow/FlowData/userdata/zimmers1/Project_SKMus73_Muscle snRNAseq_147/FeatureRankResult-2734589615408625/parameters14813225817182548634.json --output_folder
/home/flow/FlowData/userdata/zimmers1/Project_SKMus73_Muscle snRNAseq_147/FeatureRankResult-2734589615408625/out8993510355486258005 -c AICc --lognormal --useFstat --logbase 2.0 --threads 0
java: java
/opt/partek_flow/bin/gene_analysis/flow_diff_expression --input_bdb
/home/flow/FlowData/userdata/zimmers1/Project_SKMus73_Muscle snRNAseq_147/FeatureRankResult-2734589615408625/input3690637414517808576.json --model_config
/home/flow/FlowData/userdata/zimmers1/Project_SKMus73_Muscle snRNAseq_147/FeatureRankResult-2734589615408625/modelsComparisons11709814908268713496.json --attributes
/home/flow/FlowData/userdata/zimmers1/Project_SKMus73_Muscle snRNAseq_147/FeatureRankResult-2734589615408625/parameters6277911687404386154.json --output_folder
/home/flow/FlowData/userdata/zimmers1/Project_SKMus73_Muscle snRNAseq_147/FeatureRankResult-2734589615408625/out17684032807051657620 -c AICc --lognormal --useFstat --logbase 2.0 --threads 0
java: java
/opt/partek_flow/bin/gene_analysis/flow_diff_expression --input_bdb
/home/flow/FlowData/userdata/zimmers1/Project_SKMus73_Muscle snRNAseq_147/FeatureRankResult-2734589615408625/input3690637414517808576.json --model_config
/home/flow/FlowData/userdata/zimmers1/Project_SKMus73_Muscle snRNAseq_147/FeatureRankResult-2734589615408625/modelsComparisons11709814908268713496.json --attributes
/home/flow/FlowData/userdata/zimmers1/Project_SKMus73_Muscle snRNAseq_147/FeatureRankResult-2734589615408625/parameters17622061276928211841.json --output_folder
/home/flow/FlowData/userdata/zimmers1/Project_SKMus73_Muscle snRNAseq_147/FeatureRankResult-2734589615408625/out1552472969932804950 -c AICc --lognormal --useFstat --logbase 2.0 --threads 0
java: java
/opt/partek_flow/bin/gene_analysis/flow_diff_expression --input_bdb
/home/flow/FlowData/userdata/zimmers1/Project_SKMus73_Muscle snRNAseq_147/FeatureRankResult-2734589615408625/input3690637414517808576.json --model_config
/home/flow/FlowData/userdata/zimmers1/Project_SKMus73_Muscle snRNAseq_147/FeatureRankResult-2734589615408625/modelsComparisons11709814908268713496.json --attributes
/home/flow/FlowData/userdata/zimmers1/Project_SKMus73_Muscle snRNAseq_147/FeatureRankResult-2734589615408625/parameters4268559100248736539.json --output_folder
/home/flow/FlowData/userdata/zimmers1/Project_SKMus73_Muscle snRNAseq_147/FeatureRankResult-2734589615408625/out11993154543755405754 -c AICc --lognormal --useFstat --logbase 2.0 --threads 0
java: java

/opt/partek_flow/bin/gene_analysis/flow_diff_expression --input_bdb /home/flow/FlowData/userdata/zimmers1/Project_SKMus73_Muscle snRNAseq_147/FeatureRankResult-2734589615408625/input3690637414517808576.json --model_config /home/flow/FlowData/userdata/zimmers1/Project_SKMus73_Muscle snRNAseq_147/FeatureRankResult-2734589615408625/modelsComparisons11709814908268713496.json --attributes /home/flow/FlowData/userdata/zimmers1/Project_SKMus73_Muscle snRNAseq_147/FeatureRankResult-2734589615408625/parameters9283085735192267085.json --output_folder /home/flow/FlowData/userdata/zimmers1/Project_SKMus73_Muscle snRNAseq_147/FeatureRankResult-2734589615408625/out919942058740783781 -c AICc --lognormal --useFstat --logbase 2.0 --threads 0
java: java
java: java
